# A synthetic protein as efficient multitarget regulator against complement over-activation

**DOI:** 10.1101/2021.04.27.441647

**Authors:** Natalia Ruiz-Molina, Juliana Parsons, Madeleine Müller, Sebastian N.W Hoernstein, Lennard L. Bohlender, Steffen Pumple, Peter F. Zipfel, Karsten Häffner, Ralf Reski, Eva L. Decker

## Abstract

The complement system constitutes the innate defense against pathogens. Its dysregulation leads to diseases and is a critical determinant in many viral infections, e.g.COVID-19. Factor H (FH) is the main regulator of the alternative pathway of complement activation and could be a therapy to restore homeostasis. However, recombinant FH is not available. Engineered FH versions may present alternative therapeutics. Here, we designed a synthetic protein, MFHR13, as a multitarget complement regulator. It combines the dimerization and C5-regulatory domains of human FH-related protein 1 (FHR1) with the C3-regulatory and cell surface recognition domains of human FH. MFHR13 includes the FH variant I62, which we characterized to induce improved C3b binding and cofactor activity compared to the variant V62. After comparative protein structure modelling, we introduced the SCR FH_13_, which includes an *N*-glycosylation site for higher protein stability. In summary, the fusion protein MFHR13 comprises SCRs FHR1_1-2_:FH_1-4_:FH_13_:FH_19-20_. It shows an enhanced heparin binding and protects sheep erythrocytes from complement attack exhibiting 26 and 4-fold the regulatory activity of eculizumab and human FH, respectively. Furthermore, it also blocks the terminal pathway of complement activation and we demonstrate that MFHR13 and FHR1 bind to all proteins forming the membrane attack complex, which contributes to the mechanistic understanding of FHR1. We consider MFHR13 a promising candidate as a therapeutic for complement-associated diseases.

## Introduction

The complement is a fundamental part of the human immune system and constitutes the innate defense against infection agents. It consists of approximately 50 plasma and membrane-bound proteins forming a surveillance network, whose core function is the recognition and destruction of microbial invaders (1). Complement activation can occur by the classical (CP), lectin (LP) or alternative (AP) pathways, which converge at complement component C3 activation and end up in membrane attack complex (MAC) formation, triggering lysis of invading pathogens and inflammation (2).

The AP contributes to up to 80% of the overall complement activation (3) and is spontaneously activated by hydrolysis of C3 to C3(H_2_O), with C3b-like activity. C3(H_2_O), together with the factor B (FB) fragment Bb, builds the initial C3 convertase (C3(H_2_O)Bb) in fluid phase, which cleaves C3 to C3a and C3b. C3b mediates surface opsonization and amplifies complement activation, by building further C3 convertases (C3bBb) together with FB and FD. In addition the AP acts as an amplification mechanism, even when complement was activated by the CP or LP (4). As a consequence of excess C3b, C5 is activated, either by cleavage into C5a and C5b, or without proteolytic cleavage at very high densities of C3b on target surfaces, leading to C5b-like activated C5 (5). While C5b or C5b-like activated C5 bind to C6, C7, C8 and C9 leading to MAC formation (also called terminal complement complex - TCC) and cell destruction, C3a and C5a are anaphylatoxins that trigger cell recruitment and inflammation (6) (Fig. 1a).

**Figure 1.**
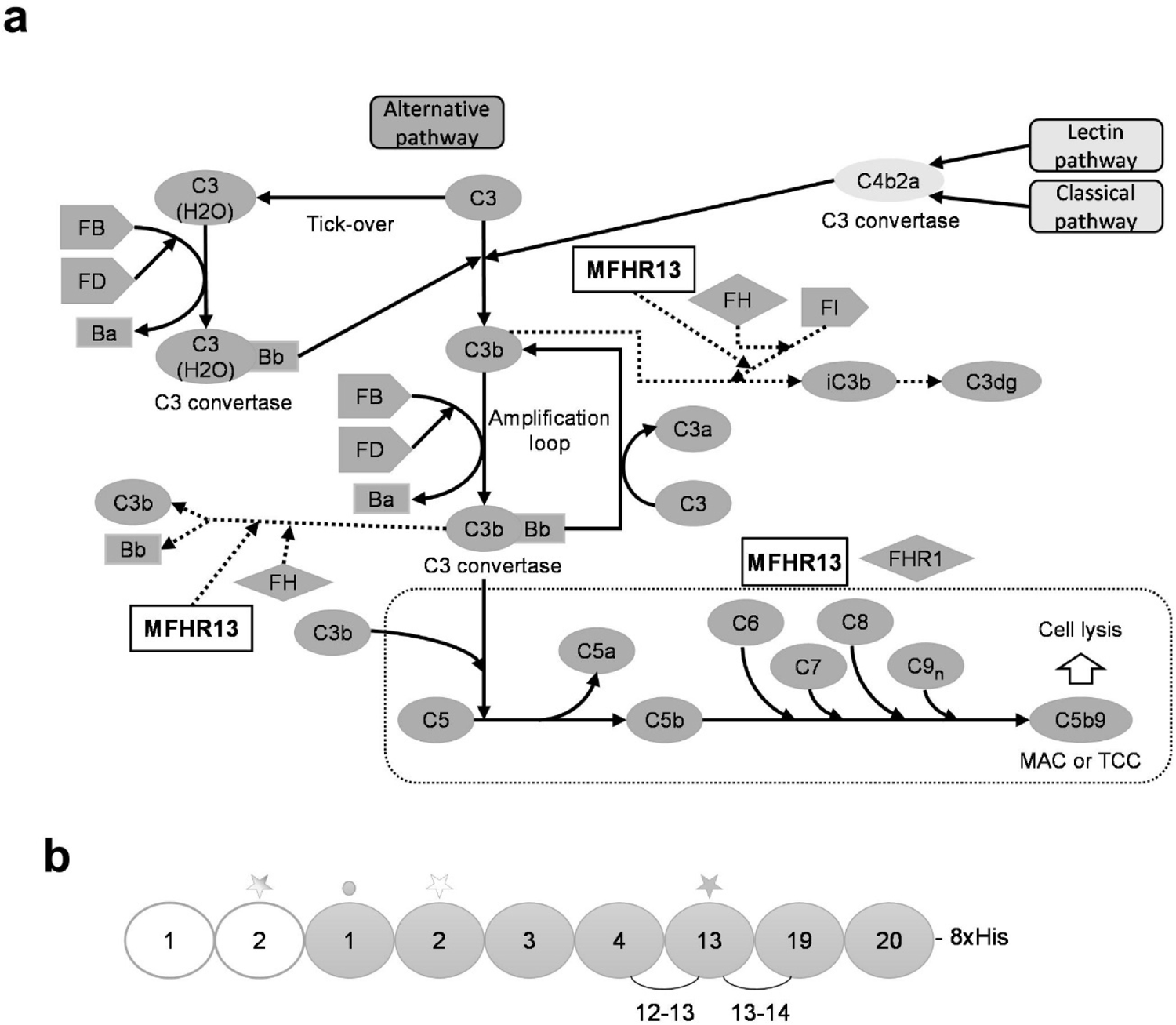
Schematic representation of complement activation and amplification by the alternative pathway (AP) and the mechanism of action of MFHR13. (**a**) MFHR13 is a regulator of the AP activation at the level of C3 and C5. The complement is activated by three pathways which converge in the formation of C3 convertases and lead to the assembly of C5b-9, called membrane attack complex (MAC), able to induce lysis and cell death. The AP is activated by spontaneous hydrolysis of C3 and acts as an amplification loop for the cleavage of C3, even when complement was activated by the classical or lectin pathways. Dotted lines and box represent sites of downregulation of AP including the sites where MFHR13 act as regulator in the cascade. (**b**) MFHR13 consists of SCRs FHR1_1-2_ (white) and 7 SCRs of FH (gray) fused by natural linkers. The gray and white stars indicate glycosylation sites and a deamidated glycosylation site, respectively. The gray dot indicates the position of the polymorphism V62I.

As complement activation can also damage intact body cells, the activation of the system is tightly controlled, especially by factor H (FH). FH is a 155 kDa glycoprotein, consisting of 20 globular domains, the short consensus repeats (SCR). The SCRs 1-4 (further named FH_1-4_) act as a cofactor for factor I-mediated proteolytic degradation of C3b to inactive iC3b (which is subsequently degraded to C3c and C3dg and finally the latter to C3d), and accelerate the dissociation of C3 convertases in serum and on host cell surfaces (7, 8). FH binds to sialic acids and glycosaminoglycans on healthy host cells, protecting them from complement attack. The cell-surface binding domains are mainly located on SCRs FH_7_ and FH_20_, while the contribution of FH_13_ is still discussed (9). FH binds also through FH_19-20_ to C3d, which contains the C3b-thioester domain (TED) that attaches to cell surfaces (10, 11).

Five factor H-related proteins (FHRs) fine-tune FH-regulatory activity. The C-terminal region of FH (FH_19-20_) is highly conserved in all FHRs, while none of them contains regions homologous to FH_1-4_ (8). Of particular interest is FHR1, a regulator of C5 activation, which inhibits the last steps of the complement cascade (terminal pathway) and the MAC formation (10, 12).

Mutations in FH, FHRs or other complement-related genes are associated with diseases such as age-related macular degeneration (AMD), atypical hemolytic uremic syndrome (aHUS), C3 glomerulopathies (C3G), and paroxysmal nocturnal hemoglobinuria (PNH) (8, 13). However, protective haplotypes have also been identified; one of them is the polymorphism FH^V62I^ (rs800292). Individuals carrying the variant I62 are less prone to complement-dysregulation diseases like aHUS, C3G, AMD or dengue hemorrhagic fever (14, 15). Furthermore, FH^I62^ showed increased binding affinity for C3b and enhanced cofactor activity (16).

Although the complement system should protect the body against viral infections, its over-activation has been associated with viral pathogenesis, e.g. in hepatitis C (17), dengue virus (18) and coronavirus infections (19, 20). Higher levels of C3a and C5a were detected in sera of patients with severe dengue (18) and deposition of complement proteins on hepatocytes was associated with liver damage in fatal cases (21). Recently, the role of complement over-activation in SARS-CoV pathogenesis has been proved. C3 deposition was found in the lungs of SARS-CoV infected mice, while C3 KO mice suffered less from respiratory dysfunction (20). Likewise, C5a accumulation and C3 deposition were observed in lung biopsy samples from COVID-19 patients (19), and enhanced activation of the alternative pathway was associated with a severe outcome of the disease (22). Furthermore, C3a and C5a induce inflammation and are important in initiating the “cytokine storm”, contributing to acute lung injury in COVID-19.

Therefore, anti-complement drugs might be an effective therapy to avoid severe inflammatory response (23, 24). Indeed, different pharmacological complement inhibitors, such as C1 esterase inhibitor (25), anti-C5 antibodies eculizumab, and ravulizumab (23, 26), C3 inhibitor AMY-101 (23) and anti-C5a antibody IFX-1 (27) (the last two still in clinical trials) have been tested to treat severe cases of COVID-19, with promising results. However, the efficacy of these treatments and the best point of the complement activation cascade to target inhibition in COVID-19 patients still needs to be assessed. Furthermore, contrary to spike proteins from a common human coronavirus (HCoV-OC43), SARS-CoV-2 spike proteins were shown to activate the complement on cell surfaces mainly through the AP. Addition of FH protected cells from spike proteins-induced complement attack (28), which suggest that complement therapeutics based on FH activity might be an important alternative.

Although huge efforts are being undertaken to develop complement therapeutics, most are still in clinical development. Anti-C5 antibodies (eculizumab and ravulizumab) are efficient complement therapeutics and the only approved drugs to treat aHUS (29, 30). However, these compounds are not as effective in many patients suffering from C3G and would not be beneficial to treat severe dengue, because they do not prevent the cleavage of C3 (18). Thus, treatments also regulating the complement at the level of C3 are necessary. In general, anti-C5 antibodies, which block the terminal pathway and complement activity, are associated with a higher susceptibility to infections and are among the most expensive pharmaceuticals in the world (31).

It is therefore essential to generate new alternative therapeutic agents to treat diseases associated with complement dysregulation using approaches to control rather than block complement activation. FH is the physiological regulator on the level of C3 and therefore, FH-based therapies could restore homeostasis. However, due to the complexity of this molecule, it is desirable to produce smaller proteins with higher overall regulatory activity. Different fusion proteins including FH active domains have been developed (32–34). The synthetic multitarget complement regulator MFHR1combines the dimerization and C5-regulatory domains of FHR1 with the C3-regulatory and cell surface recognition domains of FH to regulate the activation of the complement in the proximal and the terminal pathways. This fusion protein (FHR1_1-2_:FH_1-4_:FH_19-20_) exhibited a higher overall regulatory activity *in vitro* compared to FH or miniFH (FH_1-4_:FH_19-20_) and prevented AP deregulation in models of aHUS and C3G (32).

The moss Physcomitrella (*Physcomitrium patens*) is a suitable production host for pharmaceutically interesting complex proteins, as demonstrated by the successful completion of the clinical trial-galactosidase (Repleva AGAL; eleva GmbH) to treat Fabry disease (35). MFHR1 and FH have been produced in moss with full *in vitro* regulatory activity (36–38).

Physcomitrella’s characteristics include genetic engineering via precise gene targeting, growth as a differentiated tissue in a low-cost inorganic liquid medium, long-term genetic stability, industry-scale production in photo-bioreactors (500 L), homogenous glycosylation profile, high batch-to-batch stability and glycoengineering for improved pharmacokinetics and pharmacodynamics of the biopharmaceutical (35, 39, 40).

Here, we designed MFHR13 (FHR1_1-2_:FH_1-4_:FH_13_:FH_19-20_, Fig. 1b) as a novel multitarget complement regulator produced in the GMP-compliant moss production platform. MFHR13 includes the variant I62 of FH, which we characterized to induce a higher binding to C3b and cofactor activity. After structure assessment by *in silico* modelling, we introduced the SCR FH_13_, which includes an *N*-glycosylation site for higher protein stability (41), and contributes to increased flexibility of the molecule and cell surface recognition. MFHR13 was able to protect erythrocytes from complement attack *in vitro* much more efficiently than C5-binding antibodies, FH or MFHR1 (MFHR1^V62^). Moreover, we could demonstrate that MFHR13, as well as its parental protein FHR1, are able to bind not only C5 or C5b6, but also C6, C7, C8 and C9, providing mechanistic insights into the role of FHR1 as a regulator of the complement system. We propose MFHR13 as a promising future biopharmaceutical to treat complement-associated diseases.

## Results

The protective polymorphism V62I is located in the regulatory region of FH, which is involved in C3b-binding and cofactor activity (CA). To confirm the suitability of the I62 variant as part of an improved complement regulator, we tested the influence of this single amino acid exchange on C3b binding and CA of MFHR1^I62^ compared to MFHR1^V62^, both produced in Physcomitrella.

### MFHR1^I62^ was successfully produced in Physcomitrella

MFHR1^V62^ was obtained from the moss line P1 (IMSC no.: 40838) (38), and moss lines for the production of MFHR1^I62^ were established. For this, the *Δxt/ft* moss parental line was used (50). Recombinant proteins produced in this line display *N*-glycans lacking plant-specific fucoses and xyloses, which might trigger an immune response in patients. After transfection with pAct5-MFHR1^I62^ and selection, suspension cultures were established for all surviving plants and screened for MFHR1^I62^ production via ELISA. 70% of the lines produced MFHR1^I62^ in different concentrations (Supplementary Fig S1). Four of the best producing lines were compared regarding growth and protein productivity at shake-flask scale during 26 days (Supplementary Fig. S2). The best producing line, N-179 (IMSC no.: 40839), was scaled up to a 5 L stirred bioreactor, where a productivity of 170 µg MFHR1^I62^/g FW was achieved (Supplementary Fig. S3). For structure and activity characterization, MFHR1^I62^ was purified from 8-days-old moss tissue via nickel affinity chromatography followed by anion exchange chromatography. Approximately 500 µg MFHR1^I62^/mL were obtained after purification and concentration.

### The MFHR1^I62^ variant has higher C3b binding and cofactor activity

To analyze the influence of the polymorphism V62I on protein function, we assessed the binding of MFHR1^V62^ and MFHR1^I62^ to C3b and C5, respectively, via ELISA. As expected, there was no difference between both variants in C5 binding (P=0.9328, Fig. 2a), while MFHR1^I62^ bound to immobilized C3b significantly better than MFHR1^V62^ (P= 0.0437 Fig. 2b, Supplementary Table 2).

**Figure 2.**
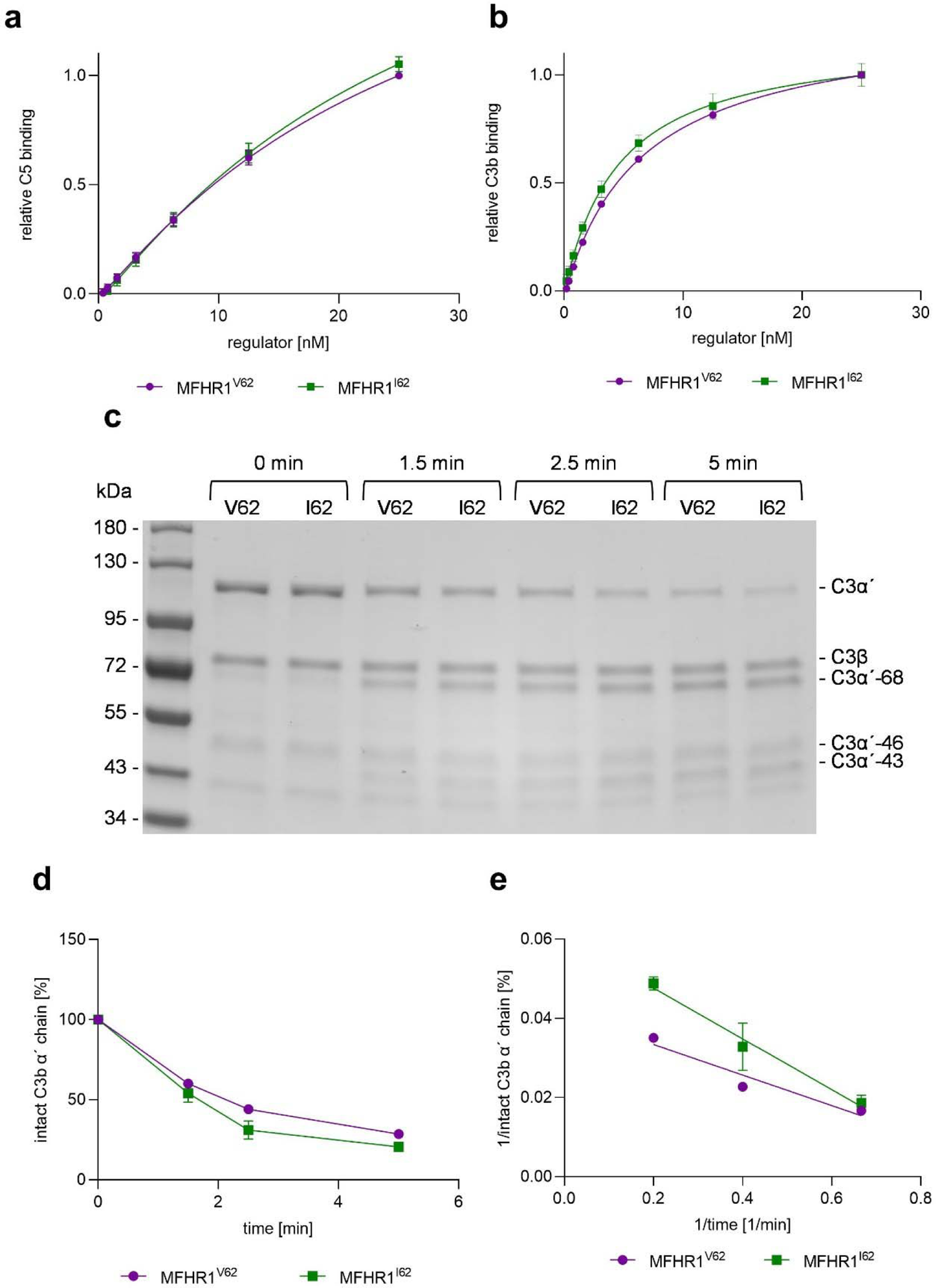
MFHR1^I62^ displays better C3b binding and cofactor activity than MFHR1^V62^. (**a**) Binding to immobilized C5 was evaluated by ELISA with increasing concentrations of MFHR1^I62^ and MFHR1^V62^. Nonlinear regression using the sigmoidal equation and logarithmic transformed data showed no significant differences between both variants binding to C5 (P=0.9328). Data represent mean values ± SD from 3 independent experiments. **(b**) Binding to immobilized C3b was evaluated by ELISA with increasing concentrations of MFHR1^I62^ and MFHR1^V62^. Nonlinear regression using the sigmoidal equation (dose response curve, 4 parameters) showed significant differences between both variants binding to C3b (P=0.0437). (**c**) Cofactor activity of FI-mediated proteolytic cleavage of C3b α‘-chain in the presence of MFHR1^I62^ or MFHR1^V62^ was visualized by SDS-PAGE and Coomassie s^α^taining. One representative experiment is shown. (**d**) Densitometric analysis of C3b cleavage. The amount of intact α’-chain was normalized for each sample with the β set to 100% intact C3b α’-chain at time zero. Two independent experiments were included in the analysis. (**e**) Double reciprocal plot of the percentage of intact C3b α analysis indicated significant differences between the MFHR1 variants (P=0.0302).

CA of both variants was measured in fluid-phase. Degradation of C3b is characterized by the cleavage of the C3b α‘-chain to fragments C3b α‘-68, −46, and −43, while C3b β-chain is not cleaved (Fig. 2c). MFHR1^I62^ displayed slightly better CA than MFHR1^V62^ (Fig. 2d). Double reciprocal plot and linear regression analysis showed significant differences between them (P=0.0302, Fig 2e). Due to the positive influence of I62 in the regulatory activity, we included this variant in MFHR13.

### Comparative protein structure modelling approach of MFHR13

To provide a potent complement biopharmaceutical, structural features that promote the efficacy of the product have to be taken into account: 1) Improving flexibility of the molecule by increasing the distance between FH_4_ and FH_19_, which will be important to enhance the biological activity of engineered versions of FH (57). 2) Improving glycosylation, since it plays an important role in the stability, half-life in the circulation, efficacy and safety of biopharmaceuticals (41). MFHR1 has two glycosylation sites. The first site in FHR1_2_ is partially occupied and the second site located in FH_4_ is deamidated and thus not glycosylated (38, 58), resulting in a high portion of non-glycosylated product. Therefore, we included an additional SCR with an *N*-glycosylation site to MFHR1^I62^ between FH_4_ and FH_19_. The glycosylation sites in FH are found in SCRs 9, 12, 13, 14, 15, 17 and 18. An interesting domain was FH_13_ due to the high amount of basic amino acid residues and its electrostatic properties, which suggests it might contain a polyanion (glycosaminoglycans or sialic acid) binding site, thus potentially increasing the host cell surface recognition.

A comparative protein structure modelling approach (Modeller 9.19) was carried out to evaluate the structure of the fusion protein FHR1_1-2_:FH_1-4_:FH_13_:FH_19-20_ and if the C3b binding sites are accessible.

To build the model we used crystal structures retrieved from the PDB database corresponding to FHR1_1-2_, FH_1-4_ +C3b, FH_12-13_, FH_19-20_, and FH_19-20_ + C3d. Furthermore, FH1114 (Model SAXS, FH_11-14_) was used to orient FH_13_, since FH_14_ shares about 30% sequence identity with FH_19_. Out of 400 models generated, we selected the one with the lowest DOPE score and assessed its quality using the web-based tools ProSa and PROCHECK. The overall quality of the model was acceptable and in the range of all available structures on PDB. According to the local model quality, where the energy is plotted as a function of residues’ position in the sequence, problematic parts of the model (characterized by positive energy values) were located in some linkers, such as the linker between FH_4_ and FH_13_. However, this is also observed in linkers of the native proteins of this family (Supplementary Fig. S4a, b). Moreover, according to the Ramachandran plot, 97.6% of amino acid residues are in favored regions, 1.3% in allowed regions and only 1.1% in outlier regions (Supplementary Fig. S4c). Our model indicates that the introduction of FH_13_ may not affect biological activity and might confer higher flexibility to the molecule (Fig. 3a). Finally, the MFHR13 model was superimposed with the structures 2WII and 2XQW, which correspond to FH_1-4_ bound to C3b and FH_19-20_ to C3dg, respectively. The natural linkers are flexible enough to allow FH_13_ to fold back over in such a way that allows FH_1-4_ and FH_19_ to interact simultaneously with one C3b molecule (Fig. 3b). Furthermore, there is a striking electropositive patch extending over one face of FH_13_ to FH_20_ (Fig. 3c), which might enhance the affinity to host cell surfaces.

**Figure 3.**
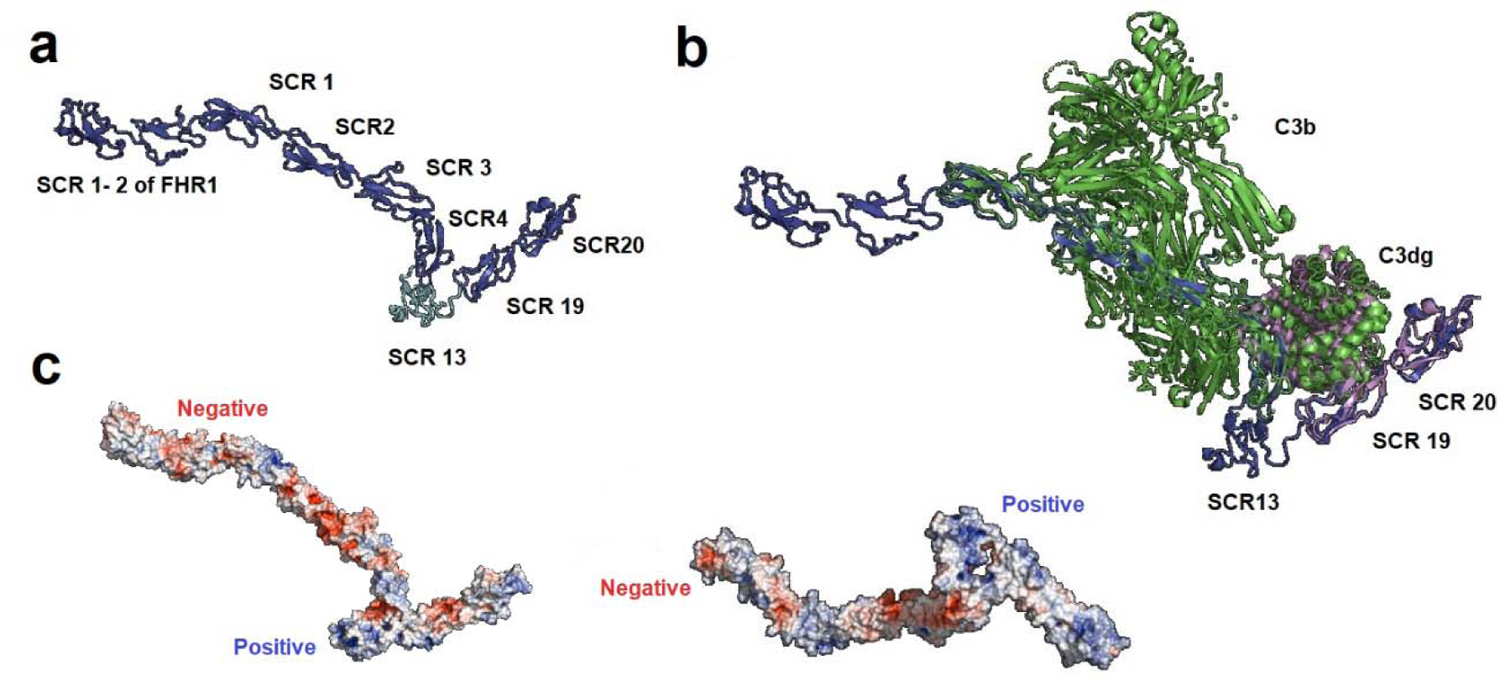
Comparative protein structure modelling of the fusion protein MFHR13 (FHR1_1-2_:FH_1-4_:FH_13_:FH_19-20_). (a) Model of MFHR13 built in Modeller 9.19. (**b**) Superimposition of the model MFHR13 (blue) with C3b (green) and C3dg (purple), respectively. Root-mean-square deviation of atomic positions (RMS) between the structure C3b: FH_1-4_ (PDB 2WII) and modelled MFHR13 is 1.345. (**c**) Electrostatic potential of MFHR13. Electronegative residues are displayed in red, electropositive in blue. The electropositive patch extending over one face of FH_13_ to FH_20_ is on the right-hand side. The electrostatic potential was determined using Adaptive Poisson-Boltzmann Solver included in PyMOL 2.0.

### Moss-produced MFHR13 is correctly synthesized and efficiently glycosylated

For the production of MFHR13 (FHR1_1-2_:FH_1-4_:FH_13_:FH_19-20_), the *Δxt/ft* moss line was transformed with the pAct5-MFHR13 construct. Selection and screening of plants producing MFHR13 was performed following the same procedure as described above for MFHR1^I62^. From 98 surviving plants, 45% produced MFHR13 in different levels as screened by ELISA (Supplementary Fig. S5).

For the best producing line N13-49 (IMSC no.: 40840) around 70 µg MFHR13/g FW were achieved in the 5 L bioreactor cultivation, which correspond to 7 mg MFHR13 after 8 days (Supplementary Fig. S6).

Approximately 200 µg MFHR13/mL were obtained after purification (Fig. 4a). The concentration was measured by ELISA and the correct ratios between MFHR13, MFHR1^I62^ and MFHR1^V62^ were further confirmed by semi-quantitative Western blot (Fig. 4b).

**Figure 4.**
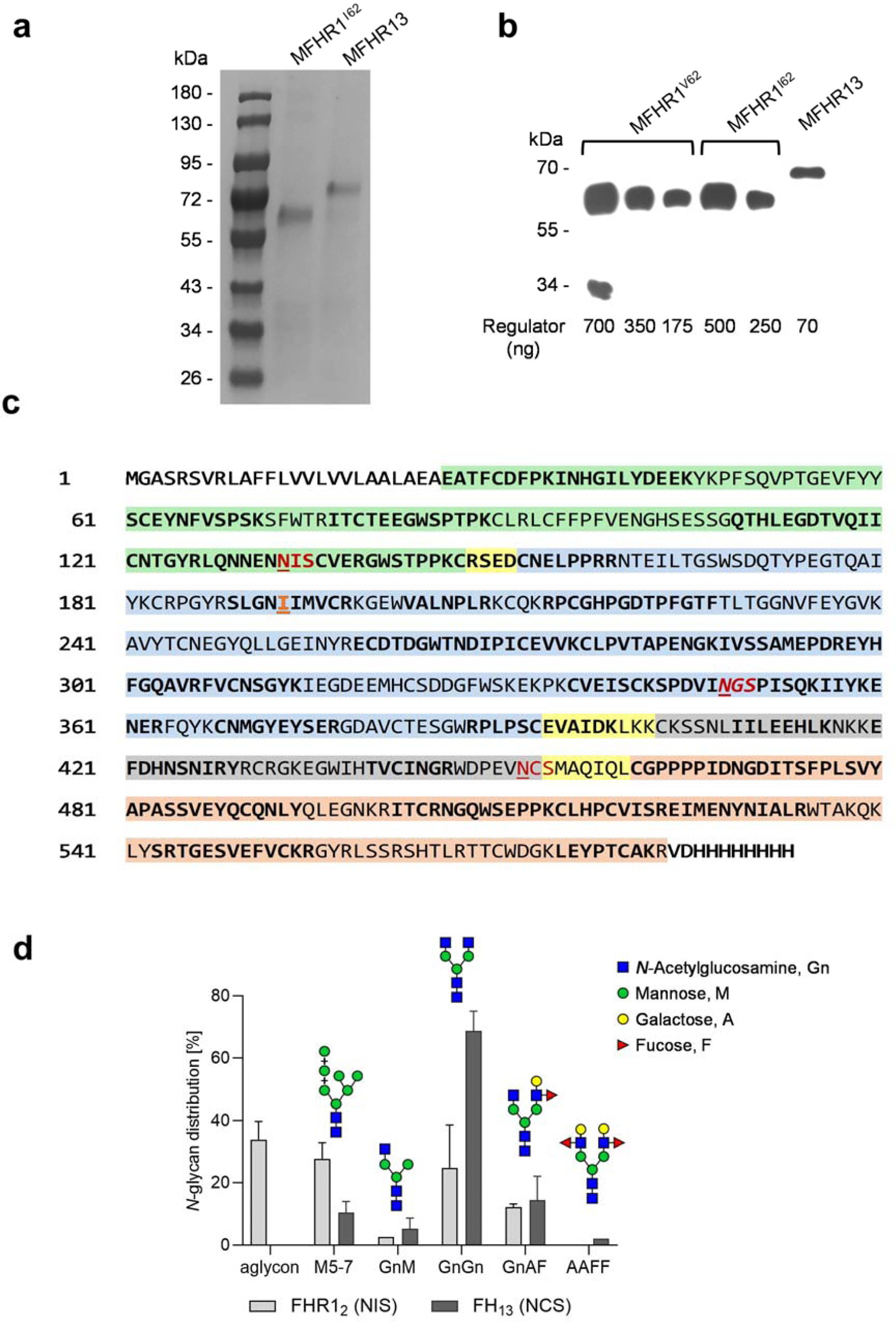
MFHR13 structure and sequence characterization. (**a)** Coomassie staining of purified MFHR1^I62^ and MFHR13 via Ni-affinity and anion exchange chromatography. Samples were separated on SDS-PAGE under reducing conditions. (**b**) Semi-quantitative Western blot of MFHR1^V62^, MFHR1^I62^ and MFHR13 under reducing conditions and detected with anti-His-tag antibodies. A calibration curve was obtained with MFHR1^V62^ and densities of the Western blot signal were compared with the concentrations obtained by ELISA (included under each lane). One representative experiment is shown. (**c)** MFHR13 sequence. FHR1_1-2_ are shown in green, FH_1-4_, FH_13_, FH_19-20_ are shown in blue, gray and light red, respectively, and the linkers are shown in light yellow. The glycosylation sites are highlighted in red and the deamidated site in italics. The peptides identified by MS are shown in bold. (**d**) Relative quantification of glycopeptides based on MS. MFHR13 was produced in stirred tank bioreactor (5 L) and extracted at day 8. NIS: glycosylation site in the FHR1_2_ in MFHR13. NCS: glycosylation site in the FH_13_ in MFHR13. Mean values from two technical replicates with standard deviation are shown.

MFHR13 migrated at approximately 70 kDa and the integrity of the protein was confirmed by Western blot and mass spectrometry (MS) (Fig. 4a-c). Via MS, MFHR13 was identified with a sequence coverage of 64%. The correct introduction of SCR13 of FH, together with its linkers, and the presence of isoleucine at position 193 were confirmed (Fig. 4c). The glycosylation site (NIS) located in the FHR1_2_ in MFHR13 was around 65% occupied with the pattern described in Fig 4d. As already described for moss-produced MFHR1 and human FH, the glycosylation site located on FH_4_ in MFHR13 was deamidated and not glycosylated. The new glycosylation site (NCS) introduced within FH_13_ was fully glycosylated, as we did not detect the unglycosylated peptide. Around 85% of all glycans comprised mature complex-type glycans (GnGn, GnAF, AAFF) and only 15% of immature glycans with terminal mannose (Fig. 4d).

### MFHR13 exhibits enhanced heparin-binding

One important feature of complement regulators is the ability to bind to polyanions such as glycosaminoglycans and sialic acids on host cell surfaces and protect them from complement attack. The ability of MFHR13 to bind to the polyanion heparin was evaluated by an ELISA-based method, and compared to MFHR1^V62^, MFHR1^I62^ and hFH, respectively. As expected, there were no differences in heparin-binding between MFHR1^V62^ and MFHR1^I62^. In contrast, MFHR13 bound to heparin twice as strong as MFHR1^V62^ and MFHR1^I62^ (P <0.0001), (Fig. 5a, Supplementary Table 3). This can be attributed to the presence of FH_13_. Heparin-bound hFH was only detected at the highest concentration tested (25 nM), indicating approximately 4-fold lower affinity to heparin than MFHR13 (Fig.5b, P<0.0001).

**Figure 5.**
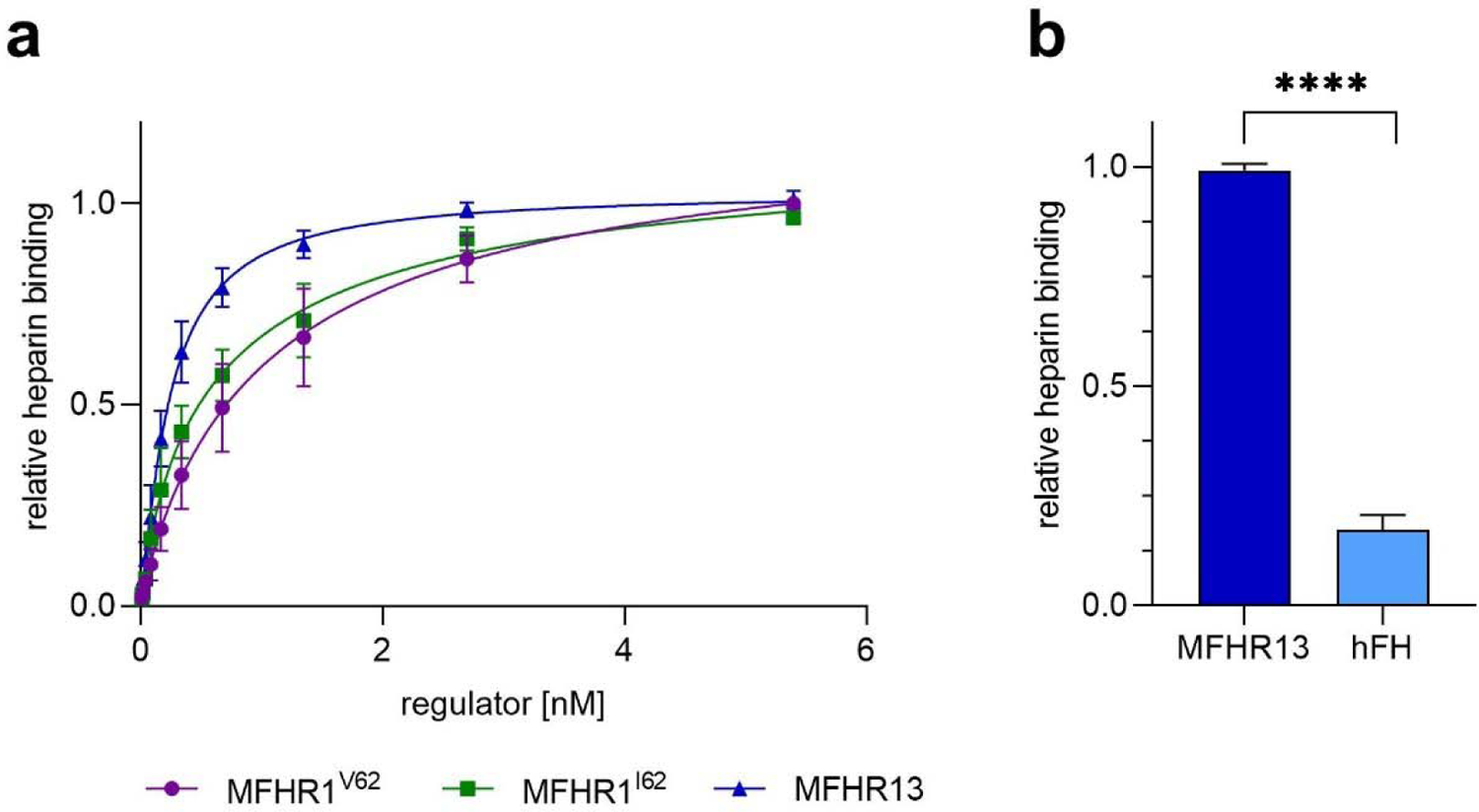
MFHR13 binds better to heparin than MFHR1^I62^, MFHR1^V62^ or hFH. (a) Heparin-binding was measured by ELISA. Nonlinear regression using the sigmoidal equation (dose response curve, 4 parameters) and comparison of IC_50_ showed significant differences between MFHR13, MFHR1^I62^ and MFHR1^V62^ binding to heparin (P<0.0001 both comparisons). Data represent mean values ± SD from 4 independent experiments. The blank (BSA 2%) was subtracted from all the samples. **(b)** MFHR13 binds significantly better to heparin than hFH (P<0.0001, unpaired t-test). The binding was measured by ELISA using 25 nM of hFH or MFHR13, respectively.

### MFHR13 regulates complement activation at the level of C3

Binding of MFHR13 to immobilized C3b was evaluated via ELISA in comparison to MFHR1^I62^, to evaluate the influence of FH_13_ on this activity. We found that both proteins bound to C3b with similar affinity (Fig. 6a; P=0.8426, Supplementary Table 4). -chain. MFHR13 showed similar MFHR13 is able to support the FI-mediated cleavage of C3b α′^I62^ fluid-phase CA in independent experiments as compared to MFHR1, however, hFH had a higher CA than both synthetic regulators. (Fig. 6b, c) (P=0.0025 and 0.0040 for MFHR13 and MFHR1^I62^, respectively).

**Figure 6.**
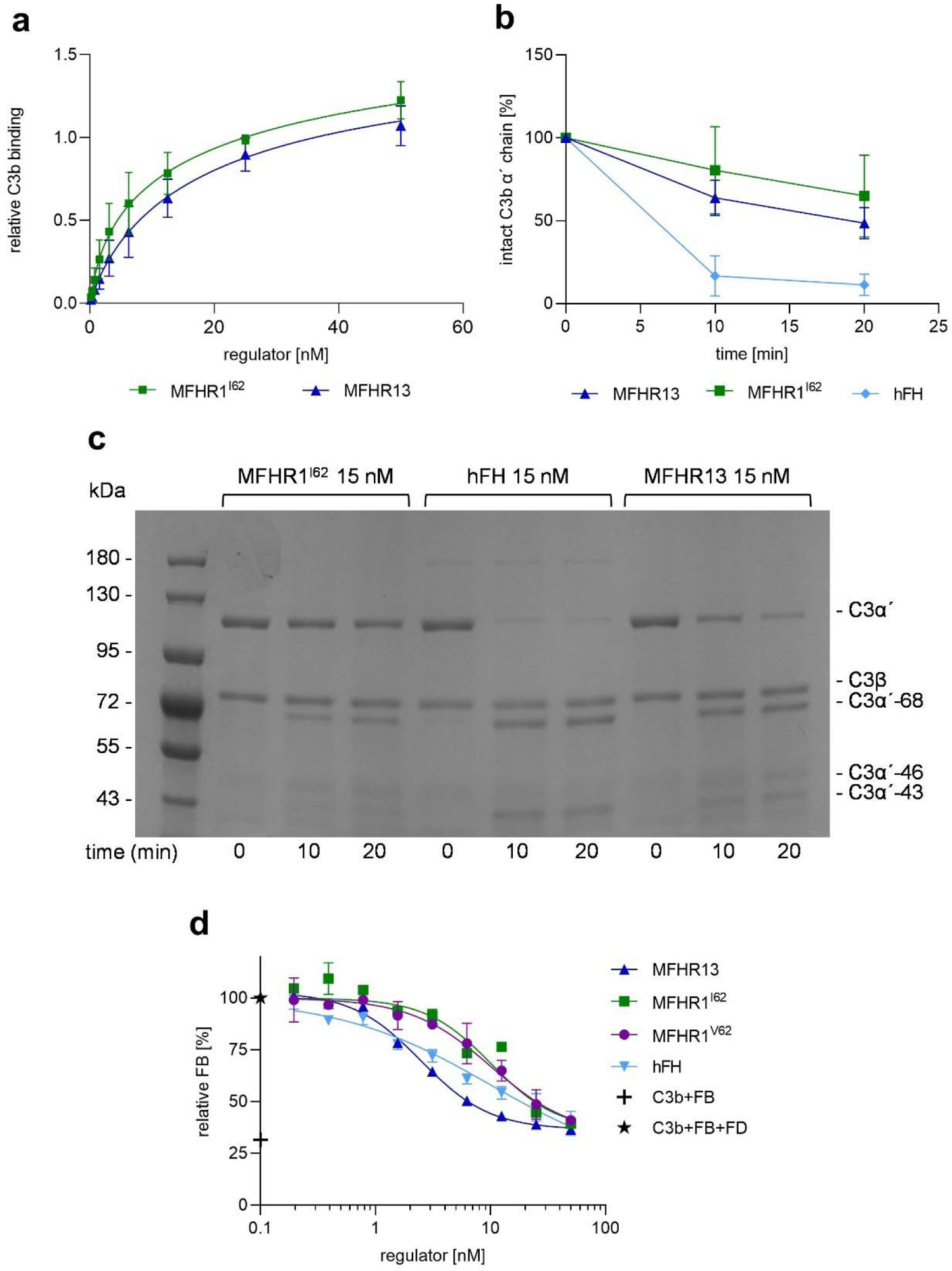
MFHR13 binds to C3b and exhibits cofactor and decay acceleration activities in fluid-phase. (a) Nonlinear regression using the sigmoidal equation (dose response curve, 4 parameters) and comparison of IC_50_ showed no significant differences between MFHR13 and MFHR^I62^ binding to C3b (P=0.8426). Data represent mean values ± SD from 5 independent experiments. **(b)** Cofactor activity of FI-mediated proteolytic cleavage of C3b ‘-chain in the presence of MFHR1^I62^, FH or MFHR13, evaluated in increasing reaction times. Densitometric analysis of C3b cleavage. The amount of intact α’-chain was normalized for each sample with the β-chain and set to 100% intact C3b α’-chain at time zero. The data represent mean value from 5 or 6 independent experiments ± SD. **(c)** One representative experiment included in b is shown. C3b, FI, and MFHR1^I62^, MFHR13 or FH were incubated for 10 and 20 min and C3b cleavage was visualized by SDS-PAGE and Coomassie staining. **(d)** MFHR13 displaced C3 convertases more efficiently than MFHR1^I62^ and MFHR1^V62^. C3bBb were assembled in C3b-coated microtiter plates with FB and FD. Complement regulators were added and the dissociation of the C3 convertases was detected by the amount of FB using polyclonal anti-FB antibodies. Absorbance of samples without regulator was set to 100% FB and samples without FD as the negative control. Nonlinear regression using the sigmoidal equation and comparison of IC_50_ indicated significant differences in DAA between MFHR13, MFHR1^V62^, and MFHR1^I62^ (P < 0.0001 for each comparison) and not significant difference between MFHR13 and hFH (P = 0.2022). The data represent mean value from 2 independent measurements ± SD.

The ability of the regulators to dissociate preformed C3 convertases, known as decay acceleration activity (DAA), was evaluated in an ELISA-based method. Increasing equimolar amounts of the regulators MFHR13, MFHR1^I62^, MFHR1^V62^, or hFH, respectively, were incubated with C3 convertase (C3bBb) and Bb fragment displacement was detected. The DAA of MFHR13, with an IC_50_ of 2.7 nM, was similar to that of hFH (IC_50_ 3.8 nM) (P=0.2022), but it significantly exceeded the activity of MFHR1^I62^ and MFHR1^V62^ (IC_50_ 7.8 and 9.4 nM respectively) (P<0.0001). (Fig. 6d, Supplementary Table 5)

### MFHR13 blocks the terminal pathway of complement

#### FHR1 and MFHR13 bind to C5, C5b6, C6, C7, C8 and C9

FHR1 and MFHR1 are able to interact with C5 (32). Binding of MFHR13 to immobilized C5 was evaluated via ELISA in comparison to MFHR1^I62^. There was no significant difference after fitting the data with 4PL nonlinear regression model (Fig. 7a; P=0.2565).

**Figure 7.**
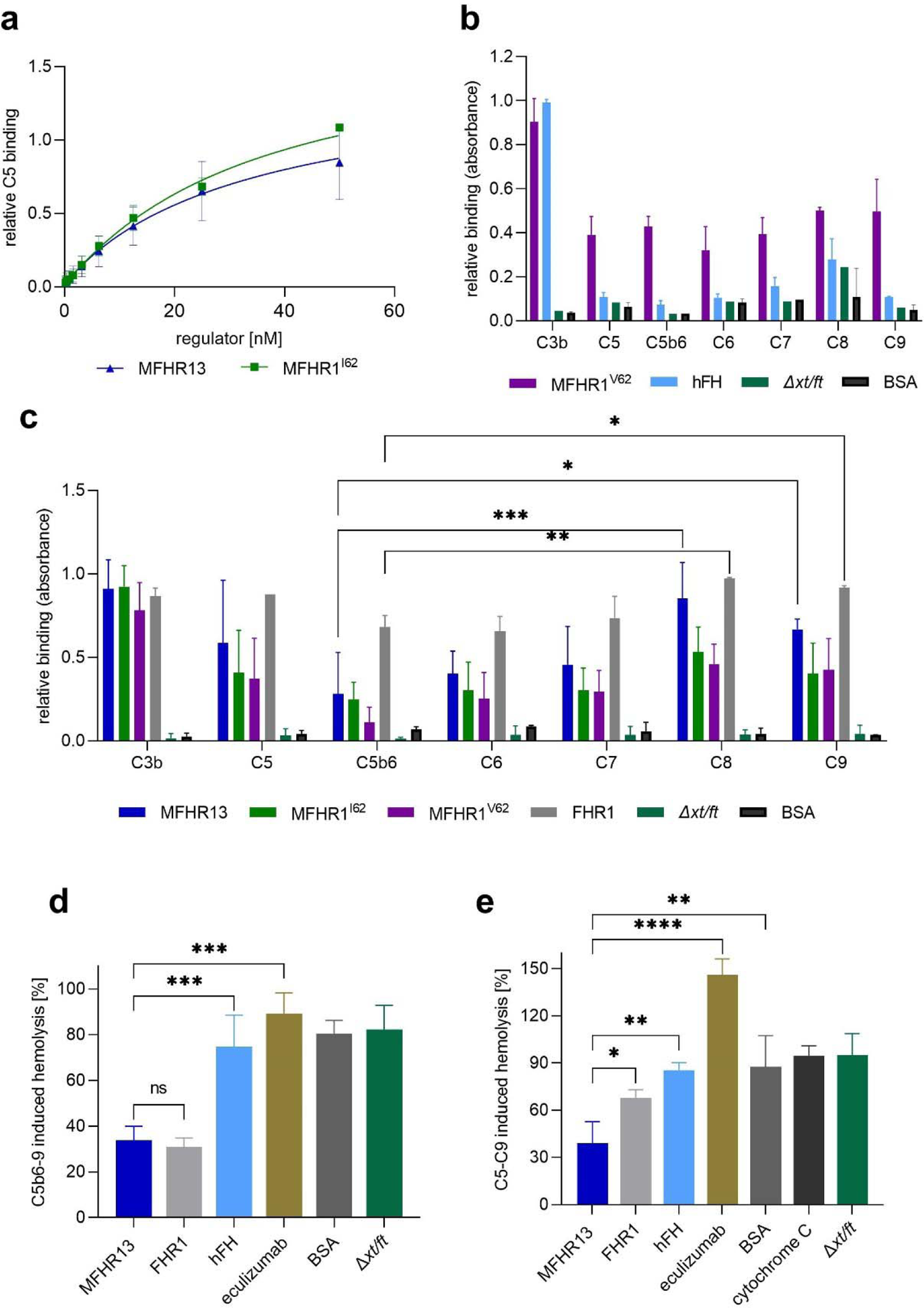
MFHR13 binds C5 and other MAC components and blocks the terminal pathway. (**a**) C5 binding was measured by ELISA. Nonlinear regression using the sigmoidal equation and comparison of IC_50_ showed no significant differences between MFHR1^I62^ and MFHR13 binding to C5 (P=0.2565). Data represent mean values ± SD from 4 or 5 independent experiments. (**b**) MFHR1 binds to the proteins involved in the MAC formation but hFH does not. MFHR1 or hFH (25nM) as well as the negative controls BSA and the purified extract from the parental line (*Δxt/ft*) were added to immobilized complement proteins in microtiter plates and detected by anti-FH polyclonal antibodies. Data represent mean values ± SD from 3 independent experiments. **(c)** MFHR13, MFHR1 and FHR1 bind to the proteins involved in MAC formation. Regulators (25 nM) were added to immobilized complement proteins and detected by anti-His-tag antibodies. Data represent mean values ± SD from 3 or 4 independent experiments. FHR1 binds significantly better to C8 and C9 than to C5b6 (P=0.0028 and P=0.0245 respectively, one-way ANOVA, Dunnett’s post hoc test). **(d)** MFHR13 and FHR1 inhibit the formation of the MAC on sheep erythrocytes significantly better than hFH (P=0.0018), eculizumab (P<0.0001), BSA (P= 0.0012), or *Δxt/ft*. Hemolysis of sheep erythrocytes was induced with C5b6, C7, C8, and C9 and determined by absorbance of released hemoglobin at 405 nm. Lysis without regulators was set to 100%. The average value of the negative controls without C5b6, and without C9 were subtracted from the positive control and samples. Data represent mean values ± SD from 3 independent experiments. (One-way ANOVA, Bonferroni post hoc test) **(e)** MFHR13 (700 nM) inhibit the formation of the MAC on C3b-opsonized sheep erythrocytes significantly better than FHR1 (P=0.0415), hFH (P=0.0018), eculizumab (P<0.0001), BSA (P=0.0012), and *Δxt/ft*. Hemolysis of C3b-coated sheep erythrocytes was induced with a mix of C5-C9 and determined by absorbance of released hemoglobin at 405 nm. Lysis without regulators was set to 100%. The negative control without C5 was subtracted from the positive control and samples. Data represent mean values ± SD from 3 independent experiments. (One-way ANOVA, Bonferroni post hoc test). **** represents P ≤ 0.0001, *** P ≤ 0.001, ** P≤0.01 and * represents P ≤ 0.05, ns (no significant difference).

Further, we tested the binding of MFHR13, the MFHR1 variants, FHR1 and hFH to C5b6 and the other proteins involved in MAC formation. Immobilized C5b6, C6, C7, C8 and C9, and C3b and C5 as positive controls, were incubated with 25 nM of MFHR13 or MFHR1-variants (8x His-tag) or 50 nM FHR1 (6x His-tag) and bound protein was detected with anti-His-tag antibodies. Due to the differences in the amount of histidine in the tag, FHR1 cannot be quantitatively compared to the moss-produced regulators. Due to the lack of His-tag, binding of hFH to the complement proteins was evaluated separately using anti-FH polyclonal antibodies and compared with MFHR1^V62^. As expected, FH bound only to C3b (Fig. 7b). In contrast, MFHR13, MFHR1 variants and FHR1 bound to all complement proteins involved in MAC formation (Fig. 7c). Here, we proved that FHR1 binds not only to C3b, C5 and C5b6 but also to C6, C7, C8 and C9, which contributes to the mechanistic understanding of FHR1 as complement regulator.

### MFHR13 prevents hemolysis by blocking MAC formation from C5b6

The C5-regulatory activity of FHR1 is still being discussed because of difficulties in separating C3 and C5-regulatory activities *in vivo*. FHR1 and MFHR1 were shown to bind C5b6 and prevent MAC formation *in vitro* (10, 32). Here, we evaluated this activity of moss-made MFHR13, using a serum-free approach and compared their activity to FHR1, hFH and eculizumab. For this, the complex C5b6 was incubated with the regulators (1 µM) followed by the addition of sheep erythrocytes and the terminal complement proteins C7, C8, and C9. Inhibition of C5b-9 formation was indirectly measured by hemolysis. The surface of sheep erythrocytes is rich in sialic acid and a good model for host-cell surfaces to test regulation of complement attack.

MFHR13, similar to FHR1, protected sheep erythrocytes from complement attack, reducing cell lysis by approximately 70%, while hFH, eculizumab, BSA and purified extract from the parental line (Δ*xt/ft*) lacked such positive effects (Fig. 7d).

The terminal pathway is initiated upon cleavage of C5 to C5b, or independently from convertases when C5 binds to densely C3b-opsonized surfaces and acquires a C5b-like conformation (5). Subsequently, MAC formation begins with the binding of C5b, or C5b-like activated C5, to C6. Here, we evaluated the ability of MFHR13 to inhibit the C5 priming on opsonized sheep erythrocytes.

Human FH did not differ significantly from the negative controls BSA or purified extract from Δ*xt/ft*, and could barely inhibit hemolysis. Under these conditions, the sample treated with eculizumab supported hemolysis, which was unexpected and differs from what was reported before when higher concentrations of complement proteins and antibody have been used (5). In contrast, MFHR13 (700 nM) reduced cell lysis by 60%, efficiently protecting sheep erythrocytes from convertase-independent C5 activation and MAC formation, while FHR1 reduced lysis by LJ 33%.

Interestingly, the hemolysis induced by C5b6-C9 was (LJ 130%) higher than the one induced by convertase-independent activation of C5, even when the concentrations of complement proteins involved in the MAC formation were almost 20 times higher in the latter assay (Supplementary Fig. S7).

### MFHR13 exhibits efficient regulatory activity on activated AP

The ability of MFHR13 to inhibit the formation of the C5b-9 complex (TCC) after activation of the AP with LPS in normal human serum (NHS) was evaluated by an ELISA-based assay and compared to MFHR1^I62^, MFHR1^V62^, hFH and the C5-blocking antibody eculizumab. MFHR13 inhibited TCC formation very efficiently, exhibiting 37-fold and 2-fold the activity of hFH and eculizumab respectively (Fig. 8a; Table 1). MFHR13 was slightly better than MFHR1^V62^ and no significant difference was observed compared to MFHR1^I62^.

**Figure 8.**
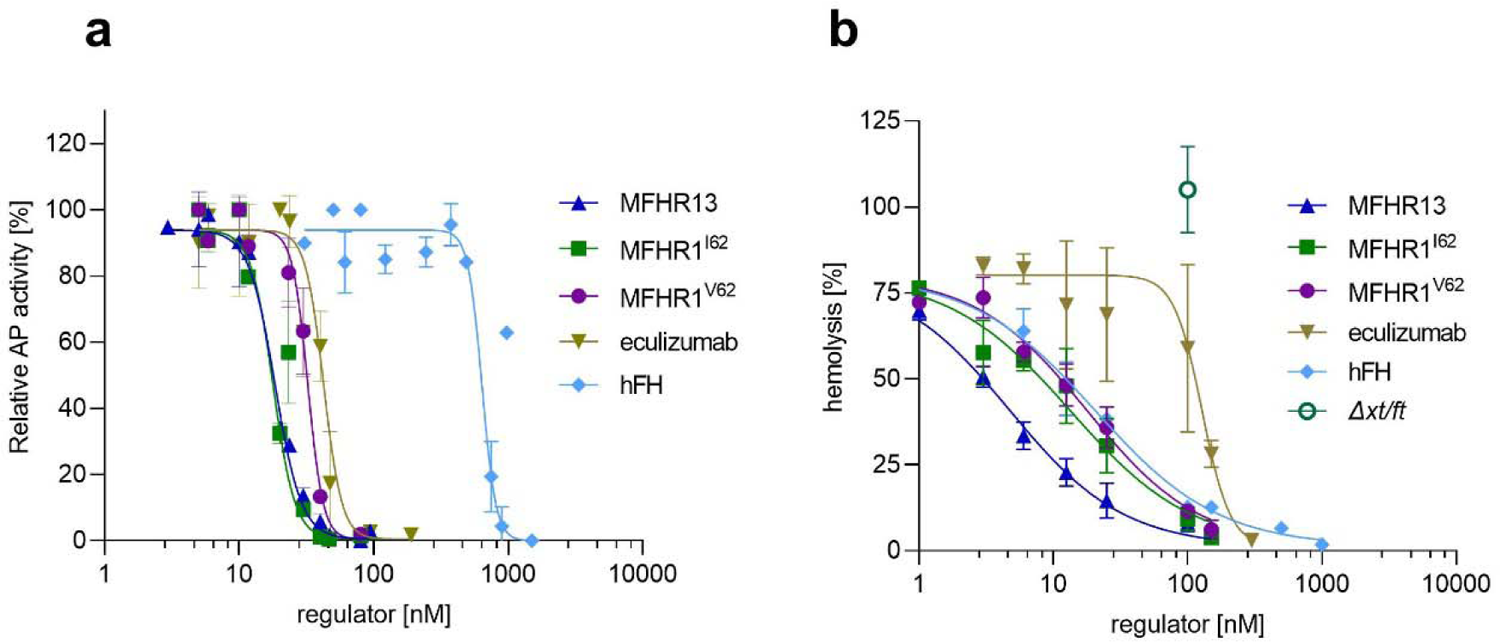
MFHR13 regulates overall AP activation and protects host-like surfaces. (a) Regulation of AP activation. AP in normal human serum was activated in LPS coated wells. The inhibition achieved by addition of regulators was evaluated indirectly by quantifying C5b-9 complex formation by an ELISA based-method. AP activity was set to 100% for wells without regulators and wells with heat inactivated serum were used as blank. Spontaneous activation was 42 ± 2.8%, measured in wells without LPS. Data represent mean values ± SD from 4 independent measurements. MFHR13 regulatory activity was significantly better than hFH (P< 0.0001, F(DFn, DFd)= 268.5 (1, 34)), and eculizumab (P<0.0001, F(DFn, DFd)= 83.38 (1, 25)). (**b**) Increasing concentrations of MFHR13 protected sheep erythrocytes from human complement attack in FH-depleted serum significantly better than MFHR1^I62^ (P=0.0003, F(DFn, Dfd) = 16.02 (1, 36)), MFHR1^V62^ (P<0.0001, F(DFn, Dfd) = 54.99 (1, 42)), hFH (P=0.0001, F(DFn, DFd))= 18.73 (1, 31)) and eculizumab (P<0.0001 F(DFn, DFd))= 111.7 (1, 33)). Hemolysis was set to 100% for wells without complement regulators and the negative control without FH-depleted serum was used as blank. Data represent mean values ± SD from 4 independent measurements. *Δxt/ft* is a purified extract from the parental moss line used as negative control. DF: degrees of freedom

**Table 1.** Overall AP regulatory activity evaluated by hemolysis and an ELISA based-method: Best fit IC_50_ calculated by 4PL nonlinear regression model. CI: Confidence Interval, R^2^: Goodness of Fit.

| Assay | MFHR13 | MFHR1<sup>I62</sup> | MFHR1<sup>V62</sup> | hFH | Eculizumab |
| --- | --- | --- | --- | --- | --- |
| **AP ELISA** |  |  |  |  |  |
| IC<sub>50</sub> (nM) | 17.8 | 19.3 | 33.1 | 665 | 38.2 |
| 95% CI | 16.9 - 18.9 | 16.1 - 22.5 | 29.4 - 37.4 | 477.2 - 893.3 | 34 - 42.9 |
| R<sup>2</sup> | 0.99 | 0.95 | 0.95 | 0.78 | 0.96 |
| **Hemolysis** |  |  |  |  |  |
| IC<sub>50</sub> (nM) | 4.9 | 14.3 | 18.4 | 21.3 | 128.8 |
| 95% CI | 3.7 - 7.2 | 10.8 - 21.7 | 14.3 - 27.2 | 14.8 - 37.0 | 115.5 - 152.6 |
| R<sup>2</sup> | 0.96 | 0.89 | 0.96 | 0.98 | 0.70 |

### MFHR13 protects sheep erythrocytes from complement-induced lysis

Finally, we tested the ability of the moss-produced regulators to protect sheep erythrocytes from complement attack. The AP was spontaneously activated by addition of MgEGTA to NHS depleted of FH. In this *in vitro*-model for host cell surfaces, MFHR1^V62^ and MFHR1^I62^ were as active as hFH, whereas MFHR13 protected erythrocytes from complement attack more efficiently than the other factors, MFHR1^V62^ (3.7-fold), MFHR1^I62^ (2.9-fold), hFH (4.2-fold), and eculizumab (26-fold) (Fig. 8b, Table 1).

## Discussion

Complement dysregulation due to mutations in complement genes, autoantibodies against complement proteins or over-activation in response to certain viral infections leads to a long list of diseases. C3G, PNH and aHUS are examples where complement is the main driver of the disease (8, 13, 59). Although the development of drugs to regulate the complement response and restore homeostasis is of huge interest, C1 esterase inhibitor and blocking antibodies against C5 are the only available so far (29). FH is a physiological regulator of the complement system, but truncated versions of it, such as mini-FH (AMY201, Amyndas) (57), have been proposed as better therapeutics because of an enhanced overall regulatory activity compared with FH. Moreover, drugs able to regulate the activation cascade at several levels might be especially interesting to increase efficiency. MFHR1 combines the regulatory domains of FHR1 and FH, and exhibits a higher regulatory activity than mini-FH (FH_1-4_:FH_19-20_) (32). Our studies led to the development of MFHR13 as a potential therapeutic for complement dysregulation.

For MFHR13 design, we first considered the polymorphism V62I. The substitution of valine by isoleucine within SCR1 of FH is located in the regulatory domain. FH^I62^ is associated with lower risk for aHUS and C3G (60). Although both amino acids are biochemically similar, and both FH variants can fold correctly, the substitution has structural consequences: The FH SCRs are tightly folded forming a compact hydrophobic core, where the residue 62 is buried, and the additional methylene group from isoleucine triggers a small conformational rearrangement. Additionally, the FH^I62^ is slightly more thermally stable and rigid than the FH^V62^ (61). We studied the influence of this amino acid exchange on the activity of the synthetic complement regulator MFHR1. Binding to immobilized C3b and cofactor activity of MFHR1^I62^ were significantly higher than that of MFHR1^V62^. Therefore, we included the more active variant I62 in MFHR13. The importance of protein glycosylation on the stability, efficacy and half-life of biopharmaceuticals in serum is well known (41). Deglycosylated proteins were shown to be inactive *in vivo* (62).

MFHR1^V62^ has a higher regulatory activity than hFH *in vitro* but was surpassed by hFH *in vivo*, which was attributed to a reduced half-life in the circulation (32). MFHR1^V62^ is only partially glycosylated, due to the nature of the *N*-glycosylation sites present in this fusion protein, which are not or only partially glycosylated in the physiological counterparts FH and FHR1, respectively (38). To support protein stability and prolong half-life in serum we aimed to include an additional *N*-glycosylation site in the new MFHR. For this purpose, we searched for glycosylated SCRs from FH, which beyond achieving glycosylation without the addition of artificial sequences would increase the distance between SCRs FH_4_ and FH_19_ in the regulator. This distance has been postulated before to be crucial for the flexibility and the activity of mini-FH (57).

We proposed the introduction of FH_13_ with its natural linkers to generate the new molecule MFHR13. FH_13_ harbors many striking characteristics: Apart from displaying an *N*-glycosylation site, it is the smallest SCR in FH with the most flexible linkers and one of the most electrostatic positive domains. It has an unusual distribution of charged groups with a net charge of + 5, similar to FH_7_ and FH_20_ (63). We hypothesized that the host cell surface recognition could be enhanced by adding this SCR. A comparative protein structure modelling approach supported our hypothesis, since FH_13_ was completely exposed with a positive electrostatic patch extending towards FH_20_. This feature may increase the affinity to cell surfaces due to electrostatic interactions and thus stabilize the cell surface-binding site in FH_20_.

The production of this complement regulator was carried out in moss bioreactors, due to the advantages offered by this system. MS analysis revealed that the FH_13_-*N*-glycosylation site was fully occupied with more than 85% mature complex-type *N*-glycans, predominantly GnGn and around 15% carrying the Lewis A epitope. Although this structure is not desired, the gene responsible for the generation of this structure was identified in moss and can be eliminated by gene targeting (64). In order to increase the half-life in the circulation, *N*-glycans still need to be terminally sialylated to avoid rapid clearance from circulation (65). Stable protein sialylation is feasible in Physcomitrella (52), although the efficiency should still be improved.

In analogy to FH, MFHR13 should protect host cells against complement attack, directing the complement regulatory activity to the cell surface by interaction with polyanions. Therefore, we tested the binding to heparin of MFHR13 compared to MFHR1^I62^, our control lacking FH_13_. MFHR13 binds significantly better to heparin than MFHR1^I62^, indicating that FH_13_ retains the function of the polyanion binding site described recently for FH_11-13_ (9). Moreover, MFHR13 binds to heparin much better than hFH.

Next, we tested the regulatory activity of MFHR13 at the level of C3. From our protein structure modelling, we expected an enhanced binding of MFHR13 to C3b due to increased flexibility of the molecule. In addition, a weak C3b binding site localized to FH_13-15_ was described recently (9). MFHR13 binds efficiently to C3b, however, we could not detect significant differences to MFHR1^I62^. This indicates that FH_13_ alone is not enough to build the C3b-binding site described for FH_13-15_.

MFHR13 displayed similar cofactor activity in fluid phase to MFHR1^I62^, while hFH outperformed both, possibly due to small steric hindrances generated by the dimerization domains present in FHR1_1-2_. MFHR13 had a similar decay acceleration activity compared to hFH and significantly higher than MFHR1^I62^ or MFHR1^V62^. Our results suggest that MFHR13 may benefit from the space between FH_4_ and FH_19_ to bind to C3b using both sites simultaneously, which could increase the decay of the convertases (C3bBb).

We demonstrated that FHR1, MFHR13 and the MFHR1 variants are able to bind not only to C5 and C5b6, but also to C6, C7, C8 and C9. The binding to C8 and C9 was significantly higher compared to C5b6, which suggest that regulation is not only mediated through C5b6-complex binding, as currently assumed (12). Our observations indicate an FHR1 activity resembling the function of vitronectin, which is a soluble regulator of the MAC (66), and thus contributes to the understanding of the mechanism of action of FHR1.

To distinguish the specific activity on C5 level from overall regulation, we evaluated the inhibition of the C5b-9 complex (MAC) formation on sheep erythrocytes in a serum-free approach using C5b6, C7, C8 and C9. We could demonstrate the activity of MFHR13 at this level of activation, which has to be attributed to the presence of the FHR1_1-2_ as described previously (10, 32).

C5 can be enzymatically activated by C3 convertases or non-enzymatically by conformational changes when bound to highly C3b- or C4b-opsonized surfaces (5). In contrast to enzymatic activation, conformational C5 activation cannot be prevented by eculizumab, explaining the residual hemolysis observed in PNH patients treated with adequate levels of this C5 inhibitor (5). As opposed to eculizumab, MFHR13 could inhibit convertase-independent activation of C5 and MAC formation on C3b-opsonized sheep erythrocytes exposed to C5-C9. This mechanism of MAC formation, however, led to a weaker hemolysis than the one induced by C5b6-C9, even at higher concentrations of complement components in the assay, and its physiologic role should be further studied.

The regulatory activity of FHR1 at the level of C5 at physiologic concentrations is controversial (33, 67), however, the physiological concentration of FHR1 itself is still under debate (68). Moreover, it is feasible that FHR1 concentrations are elevated during infection, as suggested by increased expression of the FHR1 gene in zebrafish exposed to LPS (69). Here, we used a molar excess of FHR1 and MFHR13 with respect to MAC components to inhibit C5b6-C9 or C5-C9 mediated hemolysis, but we used a concentration of FHR1 within its physiological range (0.1 - 2.7 µM) (68).

We consider that the presence of the SCRs FHR1_1-2_ with a dimerization interface represents an important feature in MFHR13, as the regulatory activity at C5 level is fundamental and coordinates with FH activity at the level of C3. This is especially the case when the complement system has been activated by a pathogen, as observed in FHR1-homolog^−/−^ mice, which exhibited higher C3a and C5a concentrations, severe inflammation and blood coagulation, leading to organ injury after LPS-induced complement activation (70).

We tested the overall regulatory activity in an ELISA-based assay, and in a cell-based assay by measuring the inhibition of hemolysis caused by the activation of the whole cascade. MFHR13 regulates the C5b-9 complex formation significantly more effectively than hFH when AP is activated by LPS in normal human serum. Although we could not demonstrate significant differences between MFHR1^I62^ and MFHR13 in this assay, they outperformed eculizumab and FH activities by a factor of 2 and 37, respectively. Furthermore, MFHR13 protected sheep erythrocytes from hemolysis caused by complement attack, exhibiting 4-, 3-, and 26-fold the regulatory activity of hFH MFHR1 versions and eculizumab, respectively. Since the surfaces of sheep erythrocytes are rich in sialic acid, we assume that the higher affinity of MFHR13 to polyanions might explain the higher local regulatory activity on cell surfaces.

SARS-CoV-2 spike proteins can bind to polyanionic glycosaminoglycans on cell surfaces and over-activate the AP, probably by displacing FH; however, exogenous FH was shown to protect cells from spike protein-induced complement attack (28). Since MFHR13 binds to polyanions very efficiently and is a potent regulator of the AP, it might represent a promising alternative for COVID-19 therapy.

In summary, we designed MFHR13, a novel synthetic multi-target complement regulator, and produced it in moss bioreactors. MFHR13 is a glycoprotein that regulates the overall complement activity around 37 times as efficient as native human FH, and twice as much as eculizumab, without blocking the complement response completely. Taken together, MFHR13 is a potential therapeutic agent for diseases such as C3G and aHUS, and might also be of interest for different viral pathogenesis where an over-activation of the complement system is involved, such as dengue and COVID-19.

### Material and Methods

#### Comparative protein structure modelling approach

To predict the 3D structure and C3b-binding capacity of a new multitarget complement regulator, MFHR13 (FHR1_1-2_:FH_1-4_:FH_13_:FH_19-20_), we used the template-based modelling implemented in Modeller 9.19 (42).

High-resolution crystal and NMR structures for 17 of 20 domains of human FH are available in the Protein Data Bank (PDB). To build the model, we used the following structures as templates (PDB accessions): 3ZD2 (FHR1_1-2_), 2WII (FH_1-4_ +C3b), 2KMS (FH_12-13_), 2G7I (FH_19-20_), 3SW0 (FH_18-20_), 2XQW (FH_19-20_ + C3d), and a SAXS model for FH_11-14_ (Accession in Small Angle Scattering Biological Data Bank (SASBDB): SASDAZ4). As the orientation between adjacent SCR domains is difficult to predict, some restrictions were added to orient the template structures appropriately.

These restrictions were obtained by measuring distances Cα -Cα between adjacent SCRs in PDB structures such as 2WII, 3SW0 using the PyMOL Molecular Graphics System, Version 2.0, Schrödinger, LLC (PyMOL 2.0). Some additional restraints between FH_4_ and FH_19_ were taken into account, after superimposing the structures 2WII and 2XQW, which correspond to FH_1-4_ with C3b and FH_19-20_ with C3d respectively.

The alignment of MFHR13 was done using also a structure for FH_11-14_, derived from SAXS information (SASDAZ4). FH_14_ was aligned with FH_19_ because they share a sequence identity of 30%, and it would allow orienting FH_13_. The module Automodel from modeler 9.19 was used to generate 100 models. The loop regions were refined using the loopmodel class, and 4 models with loop refined were built for every standard model (total: 400 models). Optimization of the objective function was done using 300 iterations with the conjugate gradient technique and molecular dynamics with simulated annealing to refine the model. The best models were chosen comparing the Discrete Optimized Protein Energy (DOPE) score. The quality of the models was evaluated using the Ramchandran Plot SAVeS Server (PROCHECK) (43) (https://services.mbi.ucla.edu/SAVES/), ProsaWEB (https://prosa.services.came.sbg.ac.at/prosa.php) (44). 3D structures were visualized with PyMol 2.0.

#### Generation of plasmid constructs

First, the vector pAct5-MFHR1^I62^, coding for MFHR1^I62^, a MFHR1 variant with isoleucine instead of valine at position 193, which corresponds to position 62 of factor H, was created. This vector was obtained via site-directed mutagenesis using the Phusion Site-Directed Mutagenesis Kit (Thermo Fisher Scientific, Waltham, MA, USA) according to the manufacturer’s instructions. The vector pAct5-MFHR1 (38)was used as template with the primers MFHR1_I62_fwd and MFHR1_I62_rev (Supplementary Table 1), exchanging a single nucleotide (GTA to ATA, V62I). In this vector the expression of MFHR1^I62^, fused to a C-terminal 8x His-tag, is driven by the 5 region, including the 5 intron, of the PpActin5 gene (45) and the cauliflower mosaic virus (CaMV) 35S terminator. For proper posttranslational modifications, the recombinant protein was targeted to the secretory pathway by the aspartic proteinase signal peptide from *P. patens*, PpAP1 (46). The hpt cassette (47) is present to select the transformed plants with hygromycin.

For the production of MFHR13 the expression construct pAct5-MFHR13, based on pAct5-MFHR1^I62^, was generated using Gibson assembly (48). For this, the sequence coding for SCR FH_13_, together with its natural flanking linkers, was amplified from plasmid pFH (36) with the primers SCR13_overlapSCR4_fwd and SCR13_overlapSCR19_rev. This fragment was inserted in pAct5-MFHR1^I62^, previously amplified with primers SCR4_overlapSCR13_rev and SCR19_overlapSCR13_fwd (Supplementary Table 1), designed to exclude the linker between FH_4_ and FH_19_ and overlapping the sequence of the linkers from FH_13_. PCRs were performed with Phusion™ High-Fidelity DNA Polymerase (Thermo Fisher Scientific). Assembled vectors were verified by sequencing.

#### Plant material, culture, transformation procedure and protein production

Physcomitrella (*Physcomitrium patens*) was cultivated as previously described (47) on Knop ME medium (49). Lines producing MFHR13 or MFHR1^I62^ were obtained by stable transformation of the Δ*xt/ ft* moss line (IMSC no.: 40828), in which the genes coding for α1,3 fucosyltransferase and the β1,2 xylosyltransferase are knocked out (50). Moss protoplast isolation, polyethylene glycol-mediated transfection (using 50 µg of linearized plasmid DNA), regeneration and selection were performed as previously described (47).

Plants surviving the selection were transferred to liquid medium and after one month of weekly propagation, moss lines were screened for the production of the proteins of interest. For this, 6-days-old moss suspension cultures were vacuum-filtrated and 30 mg fresh weight (FW) material were analyzed by ELISA. To analyze the time course of growth and production of the protein, lines were inoculated in triplicates at 200 mg/L DW and samples were taken every 3 days for 26 days.

The best MFHR1^I62^ and MFHR13 producing moss lines, respectively, were scaled up to stirred tank bioreactors (5L), operated in batch for 8-12 days with the following conditions: pH 4.5, at 22°C, aerated with 0.3 vvm air supplemented with 2% CO_2_, stirred with 500 rpm, continuous light with an intensity of 160 µmol m^-2^s^-1^ (day 0-2) and 350 µmol m^-2^s^-1^ (day 2-8). The growth medium was supplemented with 1-naphthaleneacetic acid (10 µM NAA) at day 3 according to (38).

#### Purification of moss-produced recombinant proteins

MFHR13 and MFHR1 variants (MFHR1^I62^ and MFHR1^V62^), respectively, were extracted from vacuum-filtrated plant material. For this, 4 mL binding buffer (75 mM Na_2_HPO_4_, 0.5 M NaCl, 20 mM imidazole, 0.05% Tween-20, 10% glycerol, 1% protease inhibitor (P9599, Sigma-Aldrich), pH 7.0) were added per gram FW and the suspension was disrupted with an ULTRA-TURRAX® (10,000 rpm) and simultaneous sonication (ultrasonic bath) for 10 min on ice. After two consecutive centrifugation steps (4,500 x g for 3 min and 20,000 x g for 10 min at 8°C) the supernatant was filtered through 0.22 μm PES filters (Roth).

For chromatographic purification, the filtrate was loaded onto a 1 mL HisTrap FF column, using the ÄKTA system (Cytiva) at 1 mL/min. The column was washed with 30 column volumes (CV) of binding buffer and 3% elution buffer (100%: 100 mM Na_2_HPO_4_, 0.5 M NaCl, 500 mM imidazole, 10% glycerol, pH 7.4). The protein was eluted using a stepwise gradient (6% elution buffer for 10 CV, 17% 5 CV, 27% 3 CV, 100 % 6 CV) and collected in 0.5 mL fractions. The first five fractions obtained with 100% elution buffer were pooled and diluted with Tris buffer pH 7.6 to reach 50 mM NaCl and loaded onto a 1 mL HiTrap Q HP column (Cytiva). MFHR13, MFHR1^I62^ or MFHR1^V62^ were eluted using a linear gradient (3-100%) and elution fraction containing the protein of interest (screened by ELISA and Western blot) were pooled and dialyzed against Dulbecco’s phosphate-buffered saline (DPBS) in Slide-A-Lyzer^®^ MINI Dialysis Devices, 20 K MWCO (Thermo Fisher Scientific). Proteins were concentrated by ultrafiltration using Vivaspin 2, 10 kDa MWCO (PES membrane; Sartorius).

#### Protein quantification and Immunoblotting

MFHR13 and MFHR1 variants were quantified by ELISA using a modified protocol (36): In order to quantify the fusion proteins using plasma-derived FH (hFH) standard as a reference, a polyclonal antibody against the whole FH was avoided, due to the differences in the protein structure and molecular weight between FH and the fusion proteins. Instead, a detection antibody against the FH_1-4_, domains shared by all the proteins of interest, was used.

Microtiter plates (Nunc Maxisorp, Thermo Fisher Scientific) were coated overnight at 4°C with GAU 018-03-02 (Thermo Fisher Scientific), a monoclonal antibody that recognizes FH_20_ (1:2,000 in coating buffer (1.59 g/l Na_2_CO_3_, 2.93 g/l NaHCO_3_, pH 9.6)) and blocked with sample buffer (2% BSA in TBS (Tris Buffer Saline) supplemented with 0.05% Tween-20). Samples and hFH standard (hFH, CompTech; 12.9 pM - 1.1 nM) were diluted in sample buffer and incubated for 90 min at 37°C The proteins of interest were detected by a polyclonal anti-FH_1-4_ (1:15,000 in washing buffer (1% BSA in TBS supplemented with 0.05% Tween-20) (51) and anti-rabbit coupled to horseradish peroxidase (HRP) (NA934; Cytiva, 1:5,000 in washing buffer).

The ratio between MFHR13, MFHR1^I62^ and MFHR1^V62^ concentration was also compared with semi-quantitative Western blots. SDS-PAGE on 7.5 or 10% gels, Coomassie staining and Western blot analysis were performed as described before (38).

#### Glycosylation analysis

MFHR13, purified as described above, was used for MS analysis. In brief, duplicate samples of purified MFHR13 were reduced and alkylated as previously described (52), subjected to SDS-PAGE and subsequently stained with PageBlue^®^ (Thermo Fisher Scientific). Bands corresponding to the expected size of MFHR13 were excised and distained. Digestion was performed overnight with 0.2 µg trypsin (Trypsin Gold, Promega) and 0.2 µg chymotrypsin (sequencing grade, Promega) simultaneously at 37°C. Peptides were recovered and desalted using C18 StageTips (Thermo Fisher Scientific) and measured on a QExactive Plus Orbitrap (Thermo Fisher Scientific) as described in (38). Raw data was processed with Mascot Distiller V2.7.10 and a database search was performed using Mascot server V2.7 (Matrix Science). Processed spectra from both duplicates were searched against a database containing all Physcomitrella protein models (V3.3, (53)) as well as the sequence of MFHR13 and simultaneously against a database containing sequences of known contaminants (269 entries, available on request) using a precursor mass tolerance of 5 ppm and a fragment mass tolerance 0.02 Da. As variable modifications Gln−>pyroGlu (N-term. Q) −17.026549 Da, dehydration Glu->pyroGlu (N-term. E) −18.010565 Da, oxidation +15.994915 Da (M), deamidation +0.984016 Da (N), GnGn +1298.475961 Da (N) were specified. Carbamidomethyl +57.021464 Da (C) was set as fixed modification. Search results were loaded in Scaffold5 software (Proteome Software, Inc.) and a threshold of 1% FDR at the protein level and 0.1% at the peptide level with a minimum of 2 identified peptides was specified.

Glycopeptides were identified from processed mgf files as described in (52) using a custom Perl script. Quantitative values for identified glycopeptides were obtained from the allPeptides.txt file (available on request) from a default MaxQuant (V1.6.0.16) search on the raw data. All quantitative values were normalized against the sum of all precursor intensities from each raw file.

The mass spectrometry proteomics data have been deposited to the ProteomeXchange Consortium via the PRIDE (54) partner repository with the dataset identifier PXD025471 and 10.6019/PXD025471.

#### Activity tests

### Cofactor activity (CA)

CA was measured in fluid phase. MFHR13, MFHR1^I62^, MFHR1^V62^ or hFH (15 or 75 nM) were incubated in DPBS with 444 nM C3b and 227 nM FI at 37°C. Samples (20 µL) were collected at different reaction times up to 20 min. The reaction was stopped by the addition of 7.7 µL 4x Laemmli buffer (Bio-Rad) with 3 µL 50 mM DTT (NuPAGE, Thermo Fisher Scientific). Proteolytic cleavage of C3b was assessed by visualizing the α-chain cleavage fragments α’68 and α’43 by SDS-PAGE in 7.5% gels under reducing conditions followed by Coomassie staining. The bands corresponding to the intact C3b α’-chain were quantified by densitometry (GelAnalyzer 19.1; www.gelanalyzer.com) and normalized with the corresponding C3b-β-chain. The ratio α’-chain/ β-chain at time 0 was set to 100% intact C3b α’-chain.

### Decay acceleration activity assay (DAA)

The DAA in fluid phase was performed by an ELISA-based method as previously described (37). Briefly, 250 ng C3b in PBS were immobilized on Maxisorp plates overnight at 4°C. In order to generate the C3 convertases (C3bBb), 400 ng Factor B and 25 ng Factor D (CompTech) were incubated with immobilized C3b for 2 h at 37°C. Increasing concentrations of moss-made regulators and hFH (CompTech) were added and incubated for 40 min at 37°C to measure their ability to displace preformed C3 convertases. The Bb fragments that remain bound to C3b were detected by an anti-factor B polyclonal antibody (Merck, Darmstadt, Germany), followed by HRP-conjugated rabbit anti-goat (Dako, Hamburg, Germany). The absorbance of the preformed C3 convertase without regulators was set to 100% intact C3 convertases and the C3 proconvertase (C3bB) (FB without adding FD) was included as a negative control.

### Binding to complement proteins

The ability of MFHR13 and MFHR1 variants to bind to the complement proteins was tested by ELISA. For this, 5 µg/mL of C3b, C5, C6, C7, C8, C9 or the complex C5b6 (CompTech, USA) in coating buffer were immobilized on Maxisorp plates at 4°C overnight, blocked and subsequently incubated with increasing concentrations of MFHR13, MFHR1^I62^ or MFHR1^V62^ (0.195 – 50 nM) in the case of testing C3b and C5 binding or a single concentration (25 nM or 50 nM) in the case of testing C6, C7, C8, C9 and C5b6 binding, diluted in sample buffer. Bound regulators were detected with anti-His-tag antibodies (MAB050, R&D Systems, 1:1,000 in washing buffer) and HRP-conjugated anti-mouse IgG sheep (NA931, Cytiva, 1:5,000 in washing buffer). In order to combine all independent experiments, the absorbance was normalized with the value corresponding to the highest concentration for every binding ELISA to obtain a relative binding.

Recombinant FHR1 with a C-terminal 6x His-tag (Abcam 152006) and hFH (CompTech, USA) were included as controls. However, due to the absence of His-tag in hFH, bound-hFH was detected with a polyclonal antibody (55) (1:1,000 in washing buffer), and the HRP-conjugated anti-rabbit (1:5,000 in washing buffer). It should also be considered, that the affinity of the anti-His-tag antibodies towards FHR1 compared to MFHR13, MFHR1^I62^ and MFHR1^V62^ might be different, due to the different lengths of the His-tags.

### Heparin binding

Heparin-coated microplates (Bioworld, Dublin, Ohio, USA) were used to analyze binding of the regulators to this glycosaminoglycan (GAG) analog. Bound proteins were detected with antiFH_1-4_ (1:1,000 in washing buffer) and HRP-conjugated anti-rabbit IgG from donkey (1:2,000 in washing buffer, Cytiva).

### Overall AP regulatory activity

The ELISA-based assay used to analyze the overall ability of the regulators to inhibit TCC formation after activation of the AP with lipopolysaccharides (LPS) was performed as previously described (32) with slight modifications. Briefly, increasing concentrations of MFHR13, MFHR1^I62^, MFHR1^V62^, eculizumab or hFH (0.5-100 nM) were tested and formation of C5b-9 complex was detected using a C9 neoepitope-specific antibody (aE11, Santa Cruz Biotechnology, 1:2,000 in DPBS/0.05% Tween-20), followed by HRP-conjugated anti-mouse IgG goat (NXA931, Cytiva; 1:5,000 in DPBS/0.05% Tween-20). Samples with normal human serum (NHS) and without regulators were set to 100% AP activity. Heat-inactivated NHS (56°C for 30 min) was used as a blank. A negative control to indicate spontaneous activation was included, which consisted of NHS without LPS and regulators.

The ability of the regulators to protect sheep erythrocytes from complement-mediated lysis was measured as follows: Increasing concentrations of MFHR13, MFHR1^I62^, MFHR1^V62^, eculizumab or hFH (0.3-100 nM) were incubated with 5 x 10^7^ sheep erythrocytes followed by the addition of 30% FH-depleted serum (CompTech, USA) prepared in GVB/MgEGTA buffer (0.1% gelatin, 5 mM Veronal, 145 mM NaCl, 0.025% NaN3, 5 mM MgCl_2_, 5 mM EGTA, pH 7.3). The reaction was incubated for 30 min at 37°C and stopped with GVB/EDTA (0.1% gelatin, 5 mM Veronal, 145 mM NaCl, 0.025% NaN_3_, 10 mM EDTA). The amount of hemoglobin released was measured at 405 nm. Samples without regulators were set to 100% hemolysis, samples lacking NHS were included as negative controls and the values were subtracted from all samples.

### Regulation of MAC formation on sheep erythrocytes

The inhibition of MAC formation on sheep erythrocytes was performed as previously described (10) with modifications. C5b6 (3.5 nM) was incubated with increasing amounts of the protein of interest (MFHR13, MFHR1^I62^, MFHR1^V62^, hFH, FHR1 (R&D systems) or eculizumab, 1000 nM) for 10 minutes. Then a mixture of C7 (9 nM), C8 (0.667 nM), C9 (15 nM) and 5 x 10^7^ sheep erythrocytes was added (prepared in GVB/ MgEGTA buffer) in 50 µL total volume. Hemolysis was detected after 40 minutes at 37°C by addition of GVB/EDTA buffer. The amount of hemoglobin released was measured at 405 nm. Samples without regulators were set to 100% MAC-induced lysis. BSA or purified extract from the parental line (*Δxt/ft*) were included as controls. Two negative controls without C5b6 or C9 were included, which were subtracted from all samples.

### Regulation of convertase-independent activation of C5 and MAC formation on sheep erythrocytes

The ability of the regulators to inhibit convertase-independent activation of C5 and subsequent MAC formation was tested in a hemolytic assay. For this, C3b-opsonization on erythrocytes was carried out as previously described (56) with some modifications. To achieve maximal C3b deposition without significant MAC formation, FB was partially inactivated in FH-depleted serum at 50°C for 5 min and 45 µL of this pretreated serum were added to 1 mL sheep erythrocytes (10^9^/mL in GVB/EGTA-Mg^2+^) and incubated at 37°C for 35 min. Cells were washed 4 times with DPBS and resuspended in 1 mL GVB/EDTA buffer (CompTech) to prevent formation of C3 convertases in case of residual FB.

C5 (75 nM) was preincubated for 15 min with MFHR13, FHR1, hFH or eculizumab (700 nM). BSA, cytochrome c (Sigma 2037), or purified extract from the parental line (*Δxt/ft*) were included as controls. C3b-opsonized erythrocytes (5 x 10^7^ cells) together with C6 (110 nM) were added to the regulator mix and incubated for 10 min at 37°C. Then, C7 (120 nM), C8 (70 nM), C9 (180 nM) were added to a total volume of 50 µL. After 45 min at 37°C 100 µL GVB/EDTA were added and hemolysis was detected by measuring absorbance at 405 nm. Lysis without regulators was set to 100%. The negative control without C5 was subtracted from samples and controls.

## Statistical analysis

Analyses were done with the GraphPad Prism software version 8.0 for Windows (GraphPad software, San Diego, California, USA). For experiments involving a dose-response curve, logarithmic transformed data were fitted by a four-parameter logistic (4PL) nonlinear regression model to calculate the IC_50_ and comparison of fits was carried out using the extra sum-of-squares F test with a cutoff at P=0.05.

## Acknowledgements

This work was supported by the Deutscher Akademischer Austauschdienst DAAD (to NRM), the Wissenschaftliche Gesellschaft Freiburg im Breisgau (to NRM) and the Deutsche Forschungsgemeinschaft (DFG, German Research Foundation) under Germany’s Excellence Strategy EXC-2189 (CIBSS; to RR). PFZ acknowledges support from KIdnees Iowa City USA and from the Deutsche Forschungsgemeinschaft, Collaborative Research Center SFB1192, Immune mediated Glomerular Diseases, project B6. We thank Prof. Dr. Bettina Warscheid for the possibility to use the QExactive Plus instrument, Agnes Novakovic for excellent technical assistance and Anne Katrin Prowse for proof-reading of the manuscript.

## Data availability

All data generated in this study is included in this paper and the supplementary information. The mass spectrometry proteomics data have been deposited to the ProteomeXchange Consortium via the PRIDE partner repository with the dataset identifier PXD025471 and 10.6019/PXD025471.

## Supplementary Information

**Fig. S1.**
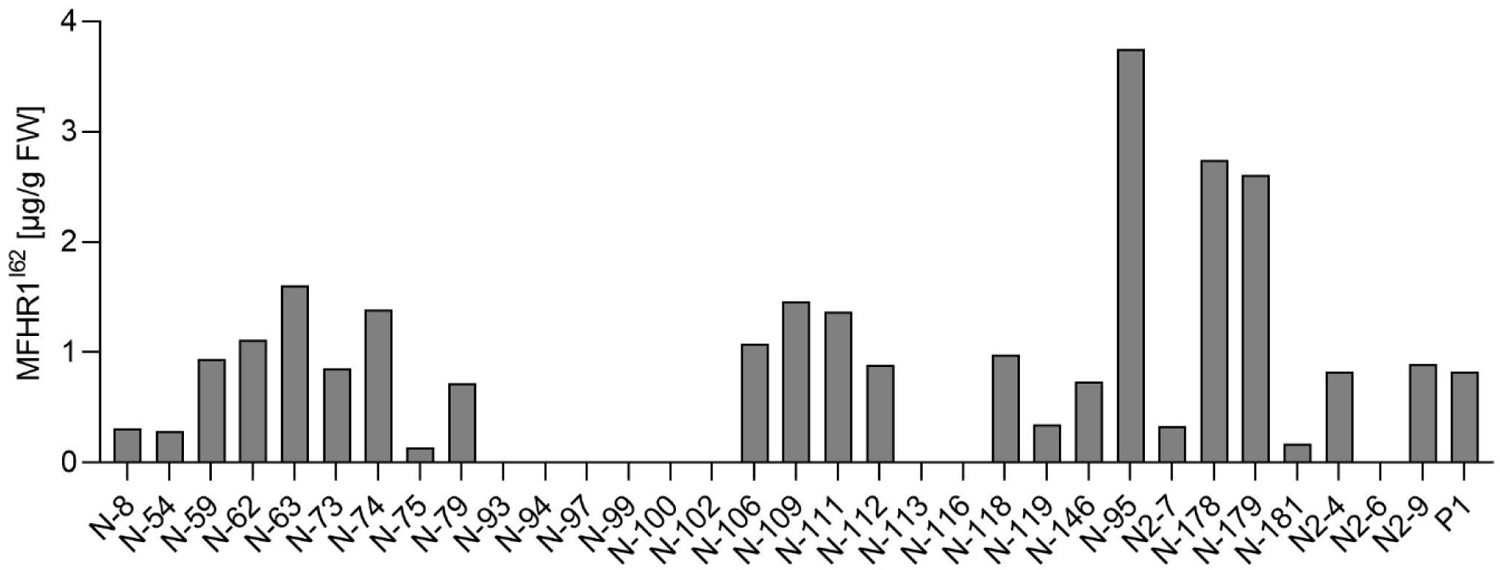
MFHR1^I62^ productivity. Plants surviving the selection were screened for productivity in suspension cultures in agitated flasks via ELISA. P1 is the MFHR1^V62^ moss producer line, which was included as an internal positive control

**Fig. S2.**
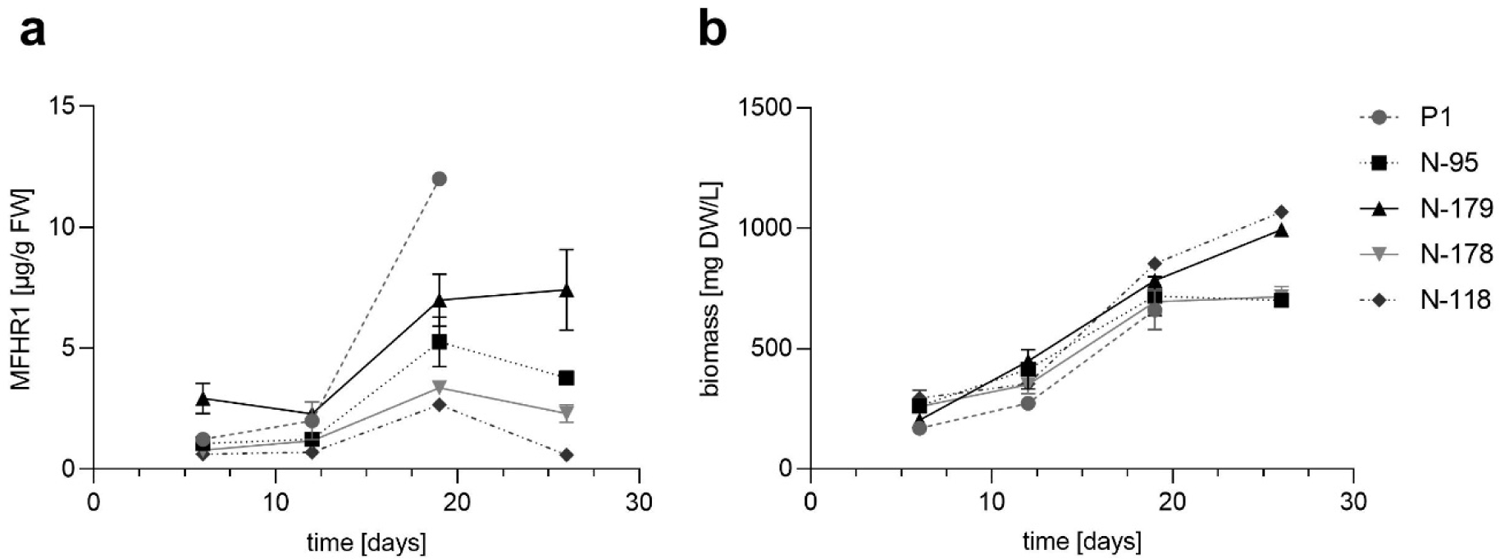
Kinetics of a) MFHR1^I62^ specific productivity and b) biomass accumulation of 4 of the best lines tested in agitated flasks.

**Fig. S3.**
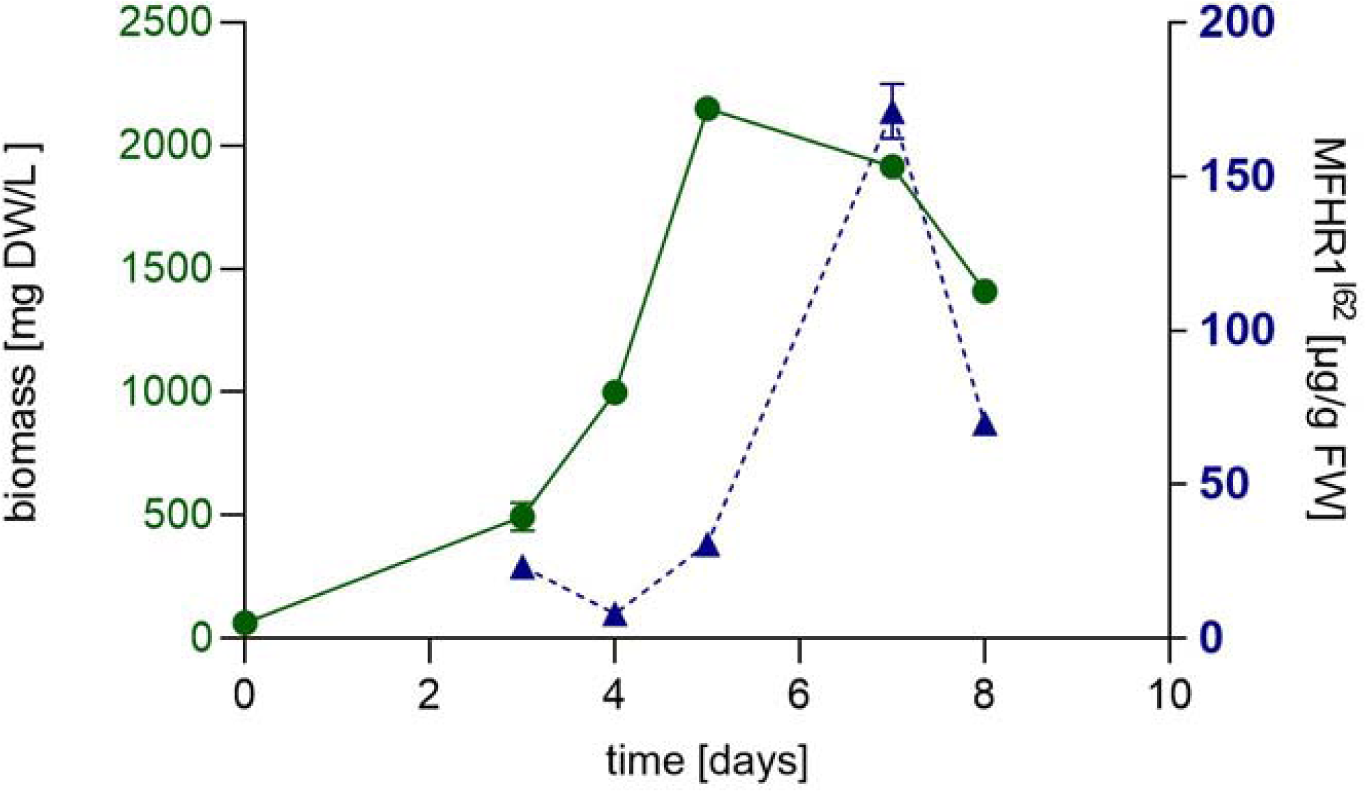
Kinetics of biomass accumulation and MFHR1^I62^ levels of plant N-179 in a 5 L stirred bioreactor.

**Fig. S4.**
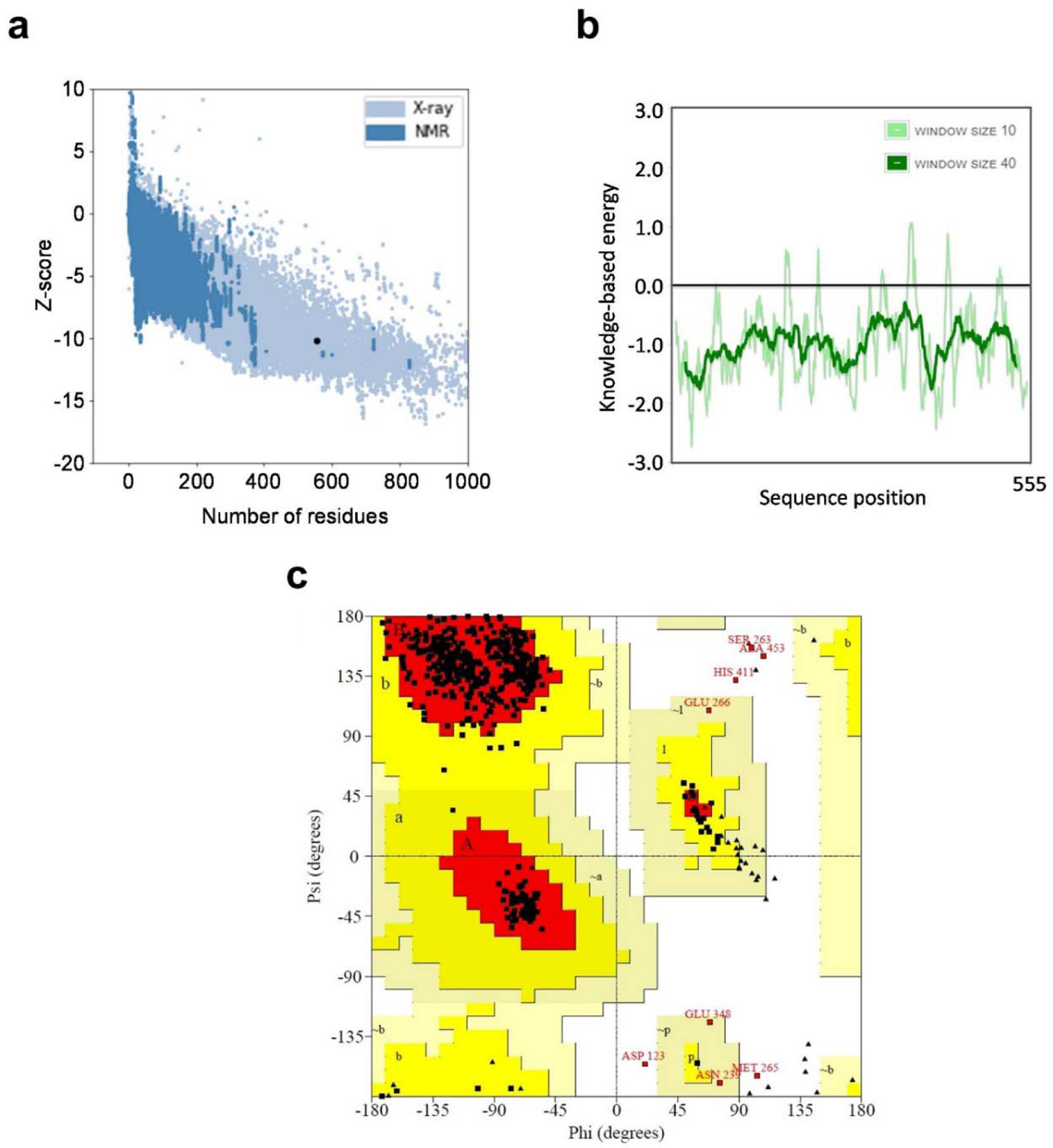
Assessment of the overall and local quality of the comparative protein structure model. (a) Overall model quality of MFHR13 obtained by ProsaWEB. Z-score: −10.07. (b) Local model quality. Diagram of energy as a function of residues sequence position. Average energy over 40 or 10 amino acid fragments was calculated by ProsaWEB. (c) Ramachandran plot analysis by PROCHECK web-based tool of the model MFHR13. The most favored regions are colored red (labeled A, B, L), allowed regions are colored dark yellow (labeled a, b, l, p), and generously allowed regions are colored in shades of light yellow (labeled LJ a, LJ b, LJ l, LJ p), while amino acids in disallowed regions are indicated as red squares. The analysis revealed that 97.6, 1.3 and 1.1% of the residues are located in favored, allowed and outlier regions, respectively.

**Fig. S5.**
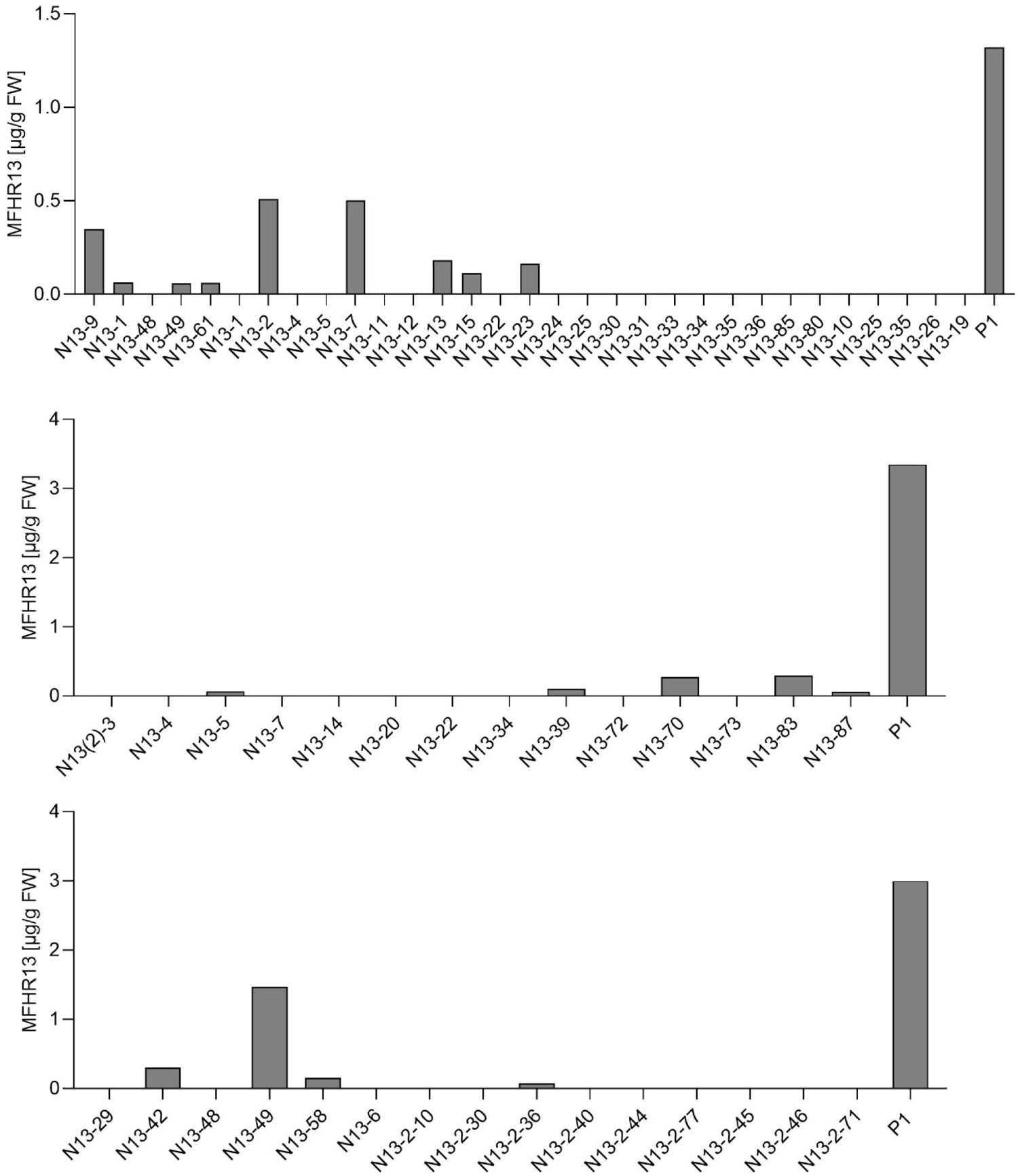
MFHR13 productivity. All plants surviving the selection were screened for productivity via ELISA in suspension cultures in agitated flasks. P1 is the MFHR1^V62^ moss producer line, which was included as an internal positive control.

**Fig. S6.**
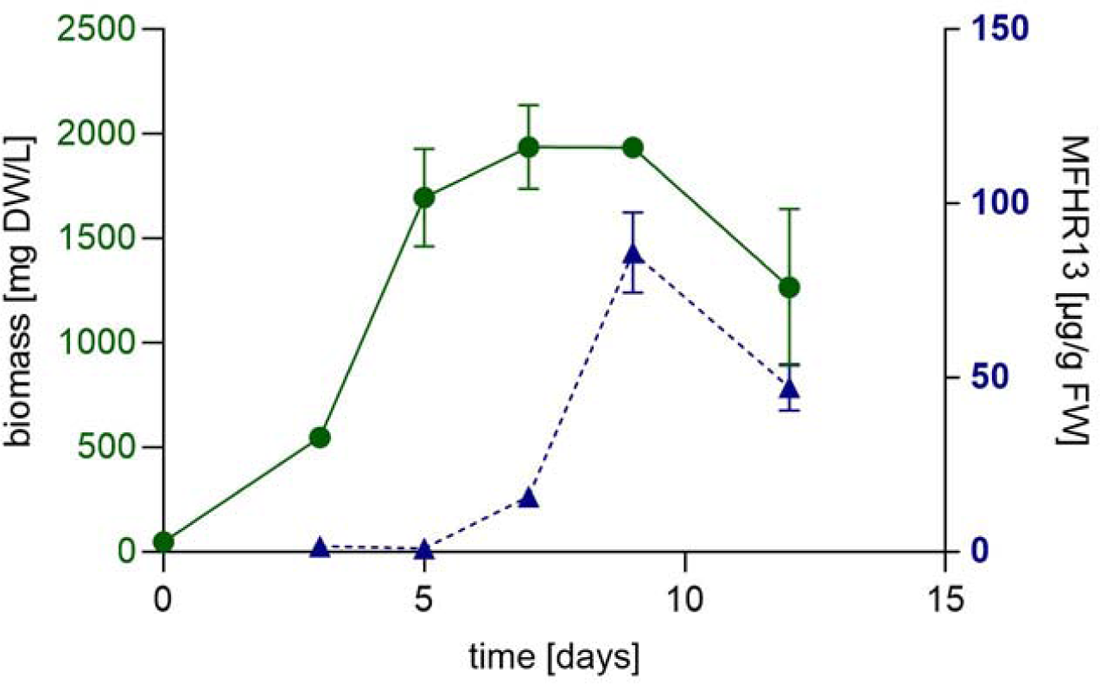
Kinetics of biomass accumulation and MFHR13 specific productivity of line N13-49 in a 5 L stirred bioreactor.

**Fig. S7.**
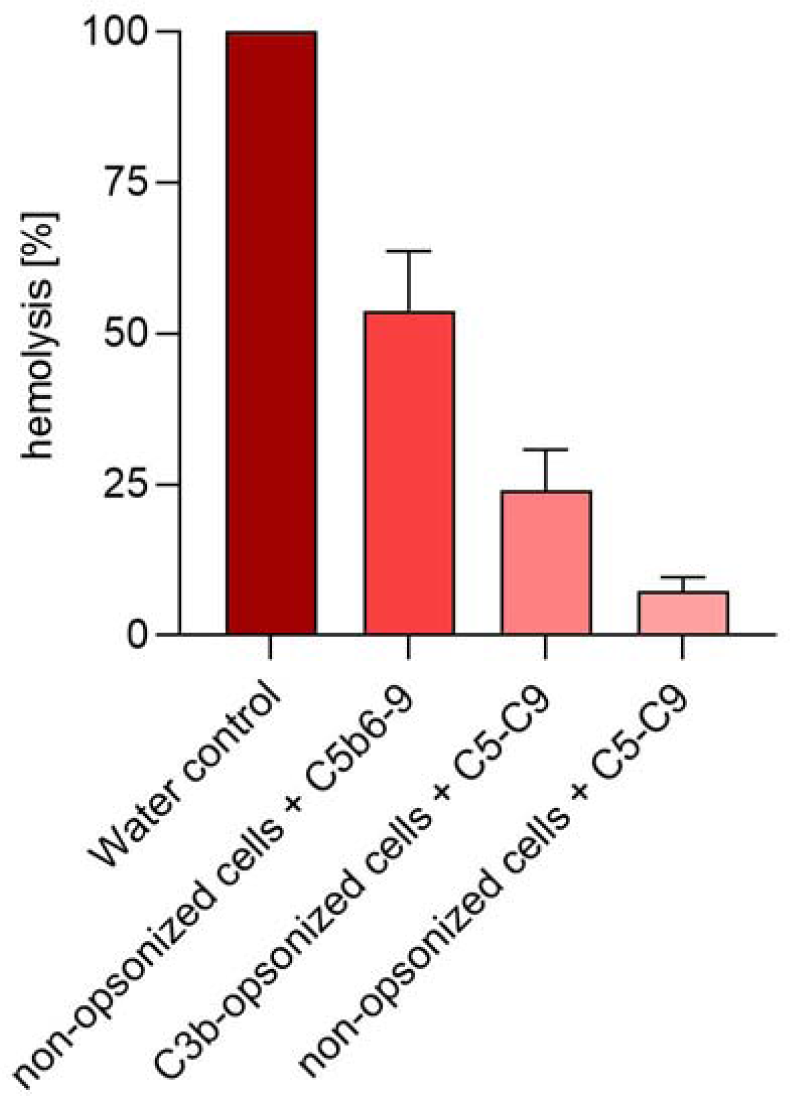
Comparison of hemolysis achieved by convertase-dependent and convertase-independent activation of C5. The extent of hemolysis achieved after activating C5 in the absence of convertases by using C3b-opsonized sheep erythrocytes exposed to C5-C9 mixture (equivalent to 20% normal human serum) is weaker than the C5b6-mediated hemolysis in non-opsonized cells. Additionally, a negative control of non-opsonized cells exposed to C5-C9 is included. Cells in water were set to 100%. Data represent mean values ± SD from 3 independent experiments.

**Table S1.**
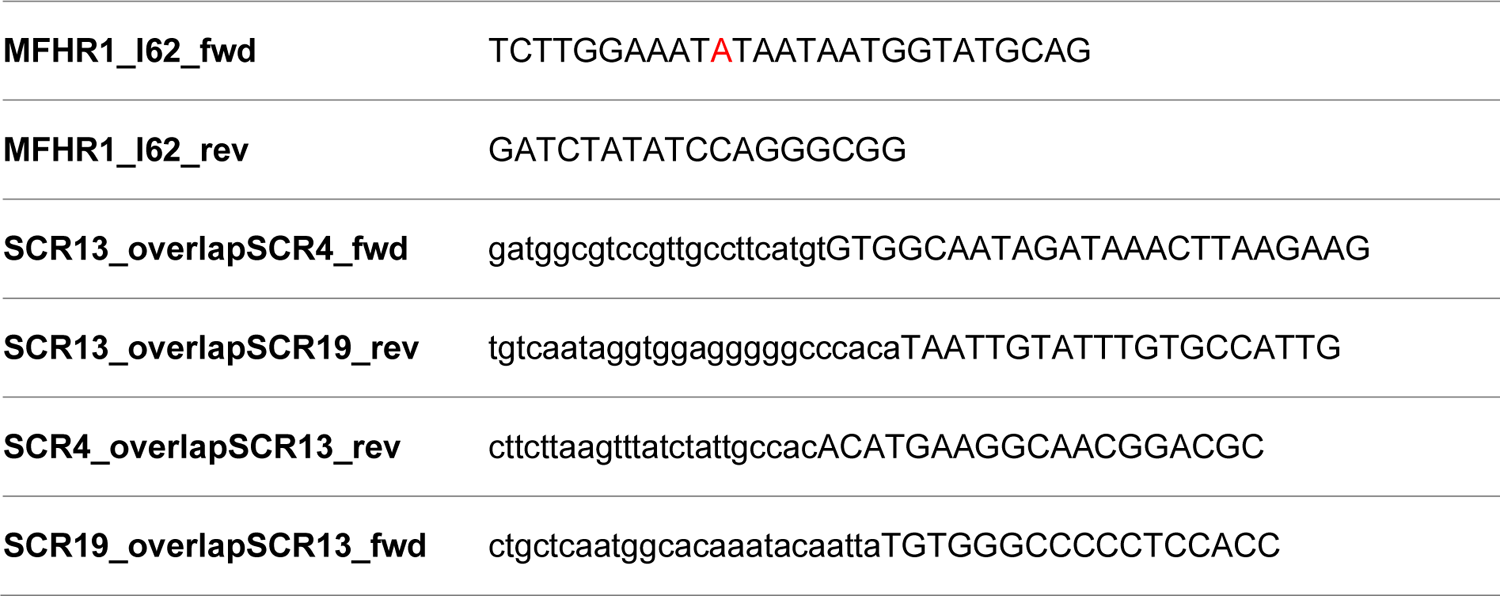
Primers used to create the expression constructs pAct5-MFHR1^I62^ and pAct5-MFHR13, by directed mutagenesis and Gibson assembly respectively. Mismatch for codon exchange is shown in red and the overhangs (overlapping regions for Gibson assembly) in lowercase.

**Table S2:**
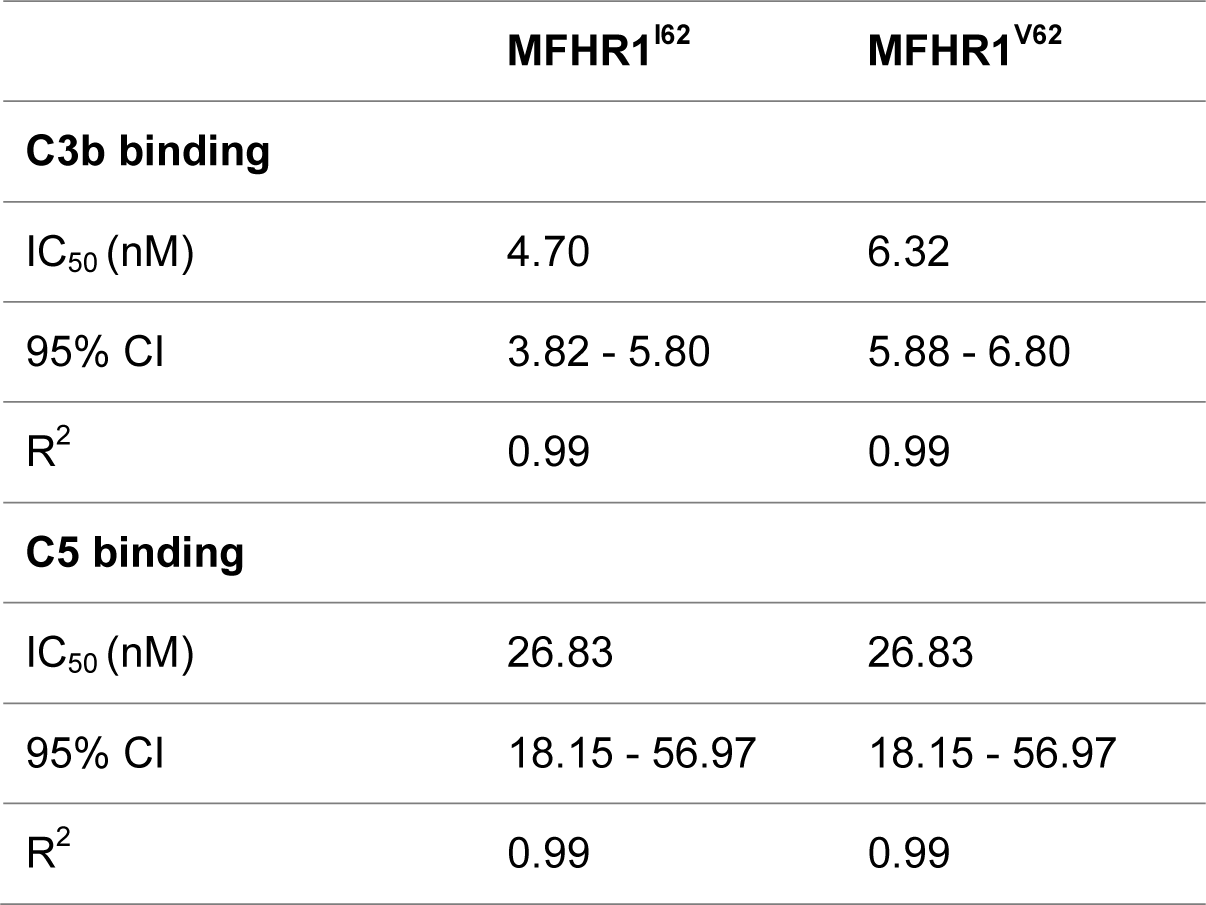
Estimated IC_50_ values for C3b and C5 binding of MFHR1^I62^ and MFHR1^V62^ by fitting the data shown in figure 2a,b with 4PL nonlinear regression model. CI: Confidence Interval, R^2^: Goodness of Fit. Comparison of fits was carried out using the extra sum-of-squares F test C3b binding: P= 0.0437, F(DFn, Dfd)= 4,338 (1, 40) C5 binding: P= 0.9328, F(DFn, Dfd)= 0.007207 (1, 40). DF= degrees of freedom

**Table S3:** Estimated IC_50_ values for heparin binding of MFHR13 and MFHR1 variants fitting the data shown in figure 5a with 4PL nonlinear regression model. CI: Confidence Interval, R^2^: Goodness of Fit. Comparison of fits was carried out using the extra sum-of-squares F test. P< 0.0001, F(DFn, Dfd)= 29.76 (2, 94). MFHR13 vs MFHR1^I62^ P< 0.0001, F(DFn, Dfd) = 23.42 (1, 66). MFHR13 vs MFHR1^V62^ P< 0.0001, F(DFn, Dfd) = 42.53 (1, 67). DF= degrees of freedom,

| **Heparin binding** | **MFHR13** | **MFHR1<sup>I62</sup>** | **MFHR1<sup>V62</sup>** |
| --- | --- | --- | --- |
| IC<sub>50</sub> (nM) | 0.24 | 0.50 | 0.76 |
| 95% CI | 0.20 - 0.28 | 0.4 - 0.65 | 0.58 - 1.17 |
| R<sup>2</sup> | 0.98 | 0.98 | 0.98 |

**Table S4.** Estimated IC_50_ values for C3b and C5 binding of MFHR1^I62^ and MFHR13 by fitting the data shown in figure 6a and 7a with 4PL nonlinear regression model. CI: Confidence Interval, R^2^: Goodness of Fit. Comparison of fits was carried out using the extra sum-of-squares F test. C3b binding P= 0.8426 F(DFn, Dfd)= 0.0397 (1, 71). C5 binding P= 0.2565 F(DFn, Dfd)= 1.309 (1, 69). DF= degrees of freedom

| **C3b binding** | **MFHR13** | **MFHR1<sup>I62</sup>** |
| --- | --- | --- |
| IC<sub>50</sub> (nM) | 16.05 | 16.05 |
| 95% CI | 8.79 - 635.1 | 8.79 - 635.1 |
| R<sup>2</sup> | 0.93 | 0.91 |
| **C5 binding** |  |  |
| IC<sub>50</sub> (nM) | 52.5 | 52.5 |
| 95% CI | Not calculated | Not calculated |
| R<sup>2</sup> | 0.86 | 0.96 |

**Table S5.** IC_50_ calculated by 4PL nonlinear regression model for decay acceleration activity (DAA), fitting the data shown in figure 6d. CI: Confidence Interval, R^2^: Goodness of Fit. Comparison of fits was carried out using the extra sum-of-squares F test. MFHR13 vs MFHR^I62^ P<0.0001 DFn, DFd = 196.6 (1, 28). MFHR13 vs MFHR^V62^ P<0.0001 DFn, DFd = 131.5 (1, 31). MFHR13 vs hFH P=0.2022 F (DFn, DFd) 1.7 (1, 30). DF= degrees of freedom

|  | **MFHR13** | **MFHR1<sup>I62</sup>** | **MFHR1<sup>V62</sup>** | **hFH** |
| --- | --- | --- | --- | --- |
| IC<sub>50</sub> (nM) | 2.70 | 7.8 | 9.45 | 3.84 |
| 95% CI | 2.2 - 3.3 | 6.2 - 9.99 | 7.7 - 11.7 | 2.95 - 5.07 |
| R<sup>2</sup> | 0.99 | 0.96 | 0.95 | 0.96 |

